# Forever Young: Structural Stability of Telomeric Guanine-Quadruplexes in Presence of Oxidative DNA Lesions

**DOI:** 10.1101/2020.11.26.399741

**Authors:** Tom Miclot, Camille Corbier, Alessio Terenzi, Cécilia Hognon, Stéphanie Grandemange, Giampaolo Barone, Antonio Monari

## Abstract

Human telomeric DNA (*h-Telo*), in G-quadruplex (G4) conformation, is characterized by a remarkable structural stability that confers it the capacity to resist to oxidative stress producing one or even clustered 8-oxoguanine lesions. We present a combined experimental/computational investigation, by using circular dichroism in aqueous solutions, cellular immunofluorescence assays and molecular dynamics (MD) simulations, that identifies the crucial role of the stability of G4s to oxidative lesions, related also to their biological role as inhibitors of telomerase, an enzyme overexpressed in most cancers associated to oxidative stress.

## Introduction

DNA G-quadruplexes (G4s) represent a non-canonical nucleic acid arrangement with remarkably different properties from the more conventional double-helical B-DNA, first described by Watson and Crick.^1^ G4s can form either in DNA and RNA, and they have been correlated to relevant biological effects also related to viral replication.^2–7^

G4s are usually emerging in guanine rich DNA regions, and their most common occurrence is based on a folded single-stranded architecture. The presence of loops connecting the different guanines also allows the formation of G4 between non-sequence contiguous nucleobases, while offering a largely increased flexibility.

From a structural point of view, G4s are constituted by a quartet of guanine bases forming planar arrangements (tetrads) and are stabilized by Hoogsteen type hydrogen bonds.^8^ These latter non-covalent interactions, have been most notably shown to present a high degree of cooperativity, also justifying the extremely high rigidity of the G4 core.^9–11^

The formation of the tetrad arrangement is accompanied by the accumulation of a quite important negative charge, that could lead to electrostatic repulsion in the center of the quartet, compromising the global stability. For this reason, the central channel is stabilized by the presence of a cation, which leads to a weak electrostatic interaction with the 6-oxygen atoms of the guanines in the tetrad,^12–14^ usually monovalent alkaline metal ions such as K^+^ and Na^+^.^13^ Furthermore, low hydration and crowded environments, such as those found in intracellular conditions, are suitable to increase the G4s stability.^15^ Indeed, G4s have been recognized as particularly stable and resistant in various conditions. These also include, albeit non exclusively, their capacity to resist to the cleavage by nuclease or the much higher thermal stability compared to other nucleic acid arrangements.

Despite the stability and rigidity of the core structure, G4s can exist in different conformation depending on the relative sugar orientation. Conventionally, this gives rise to the so-called parallel, antiparallel, and hybrid structures,^16,17^ obtained by the relative orientations of the backbone and loops connecting the tetrads. The equilibrium between the conformers is also highly sensitive to the environmental conditions and may change drastically depending on the central cation, or on the presence of crowding agents.^18^ Recently, we have shown that proper use of a strategy relying on the concomitant use of molecular simulations and spectroscopic techniques, such as electronic circular dichroism (ECD), allows to properly characterize the specific signatures and unequivocally identify their structures.^19,20^

Although, the presence of G4s in cellular, either cytoplasmatic or nuclear compartments, has been confirmed only rather recently, their biological functions are various and crucial. For instance, they are involved in chromatin remodeling, regulation of replication and gene expression and have been associated with genomic instability, genetic diseases and cancer.^21^ Indeed, G4s arrangements are also present in gene promoting regions, allowing a transcriptional regulation of the corresponding gene, as it has been well described for the oncogene *c-myc*. One of the most crucial function of G4s is also to protect the telomeric ends of the chromosomes, that comprise guanine-rich single-stranded regions. In this context G4s also act as efficient inhibitors of the telomerase,^22,23^ the enzyme controlling telomers length during replication. Since the progressive shortening of telomeres is related to the triggering of cellular senescence and death pathways, its deregulation can be related to carcinogenesis, and in particular to the “immortality” phenotype of cancer cells. As a consequence, G4 stabilizers are nowadays widely considered as potential anticancer agents and some of them are presently in clinical trial.^24–27^ More generally, maintaining of the G4s structural stability is essential to avoid triggering senescence of the cells and of the organism, especially in presence of external stress conditions.

As well as other DNA structures, G4s may be subjected to photolesions or oxidative damage that can occur as a consequence of oxidative stress. Indeed, G4s may even be regarded as hot-spots for oxidatively induced lesions since guanine is the most sensitive nucleotide to reactive oxygen species (ROS), and to oxidation in general, since, its reduction potential is the lowest among all the DNA bases.^28^ Guanine oxidative products cover a quite large chemical space,^29,30^ but 8-oxoguanine (8-oxoG) is by far the most common and hence the most characterized lesion.^31–33^ 8-oxoG may be produced from a direct one electron reaction of hydroxyl radical (OH•) on the C8 atom of a guanine, followed by an electron transfer to O_2_ and deprotonation.^30^ 8-oxoG is also produced via DNA photosensitization through the intermediate of singlet oxygen (^1^O_2_).^33,34^ Although the structure of 8-oxoG is very close to guanine, it has different physico-chemical properties, for example, its steric clash is higher. It has also been reported that 8-oxoG/C DNA strands have a consistently different hydration environment compared to G/C base pairs in normal DNA.^35^ Hence, the introduction of 8-oxoGs lesions may have a significant impact on the global structure of DNA.

If the impact of DNA lesions in G4s is less widely analyzed in comparison to that in canonical DNA, it has recently emerged that damaged G4s could lead to crucial physico-chemical or even biological outcomes that deserves attention. As a matter of fact, Markovitsi’s group has recently pointed out that a guanidinium cation, i.e. an intermediate in oxidative lesions pathways, has a much longer life-time in G4s than in canonical DNA.^36^ Furthermore, Burrows’s group has on the one side identified the presence of 8-oxoG in DNA G4s; and on the other side pointed out the specific interaction between the damaged strand and the repair protein machinery.^37^ Importantly, it has emerged that the interaction with base excision repair (BER) components may lead to a complex signaling ultimately resulting in the modulation of gene expression, and hence in epigenetic regulation. Other important biological effects have been correlated to the introduction of oxidative lesions inside the G4s sequence, among which the increase of transcription and telomerase activity.^12,38,39^ Moreover, the accumulation of oxidative lesions play an important role in both neurodegeneration and carcinogenesis,^40,41^ and hence the role of G4s as a promising target for new anticancer drugs should be considered.^25,38,42^

In a previous work we have analyzed by molecular modeling and simulation the effects of the presence of apurinic/apyrimidinic (AP) sites on the structure of *h-Telo* G4s.^43^ We have pointed out that the G4 stability strongly depends on the position of the AP in the DNA sequence. While usually the damaged G4 structure has been shown to rearrange to maintain the global folded conformation, some cases can be evidenced in which the quadruplex structure is totally unfolded. The unfolding is usually correlated with the disruption of the central leaflet and the concomitant release of both central cations.

Differently from the case of AP, 8-oxoGs in G4s usually leads to conformational changes thanks to the recruitment of guanines from the peripheral loop sequence,^44,45^ while the specific position of the damage seems to still be critical in dictate the seriousness of the unfolding.^46^ The impact of 8-oxoG lesions into G4s can be reduced by replacing guanine with xanthine,^47^ or by using a pyrene-modified guanine tract.^48^

In the present study we aim to provide a full and systematic analysis on G4s DNA damage,^43^ by investigating the structural and biological effect of single and double 8-oxoGs lesions. To this aim, we combined molecular modeling and simulation studies with electronic circular dichroism (ECD) spectroscopy of G4s exposed to increasing concentration of hydrogen peroxide (H_2_O_2_). Furthermore, we performed cellular biology assays to quantify and localize G4s and 8-oxoG via immunofluorescence in H_2_O_2_ treated cell lines. As a note, we may recall that in biological environment telomeric G4s may adopt parallel or hybrid conformation, and a precise disentangling of the conformational space is quite complicated. For this reason, while most of the MD simulations involve parallel strands, we have also checked the results for hybrid configurations. In the same spirit, we also repeated the ECD determination for both conformations obtaining coherent results.

## Results and Discussion

### Molecular Dynamics Simulations

We ran 17 simulations, with two replicas, of damaged parallel G4s structures in different orientations (Figure 1). As previously mentioned, some control was performed on hybrid structures yielding equivalent results as reported in SI.

**Figure 1.**
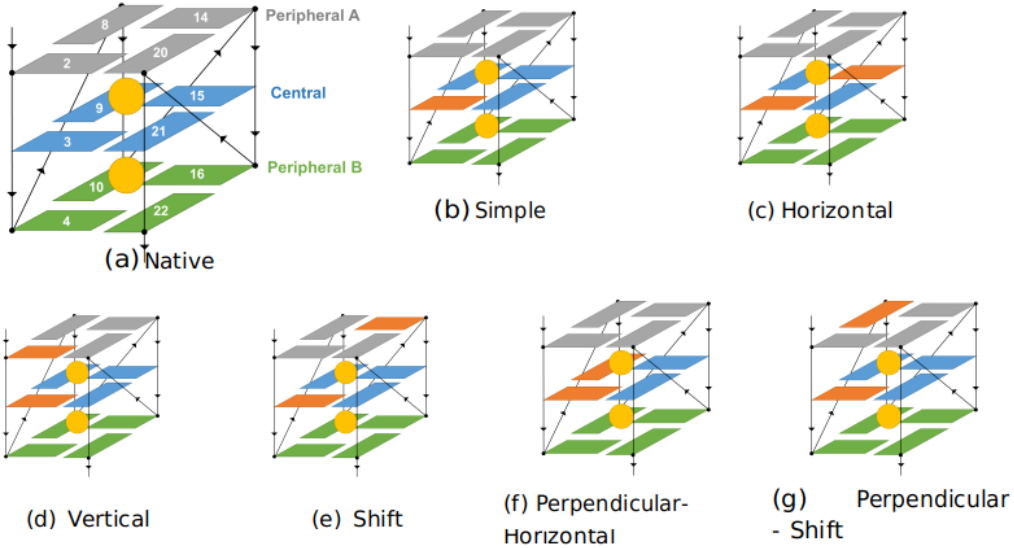
Position of the lesions in the native G4 studied in the present work and the relative orientation of the double lesions.

As also shown in Table 1, the introduction of a single 8-oxoG lesions essentially induces torsions of the quadruplex structure and the backbone, that is related to the expulsion of one central K^+^ and eventually to the disruption of one tetrad. Conversely, double 8-oxoG lesions simulations mostly totally unfold the arrangement. As a general tendency, MD show that the G4 DNA structure is globally preserved when damaged with only one 8-oxoG. This effect is also confirmed by the analysis of the root mean square deviation (RMSD) and by some particular snapshots extracted along the MD trajectory. Results are reported in Figure 2 for some representative arrangements and in the SI for the totality of the trajectories.

**Figure 2:**
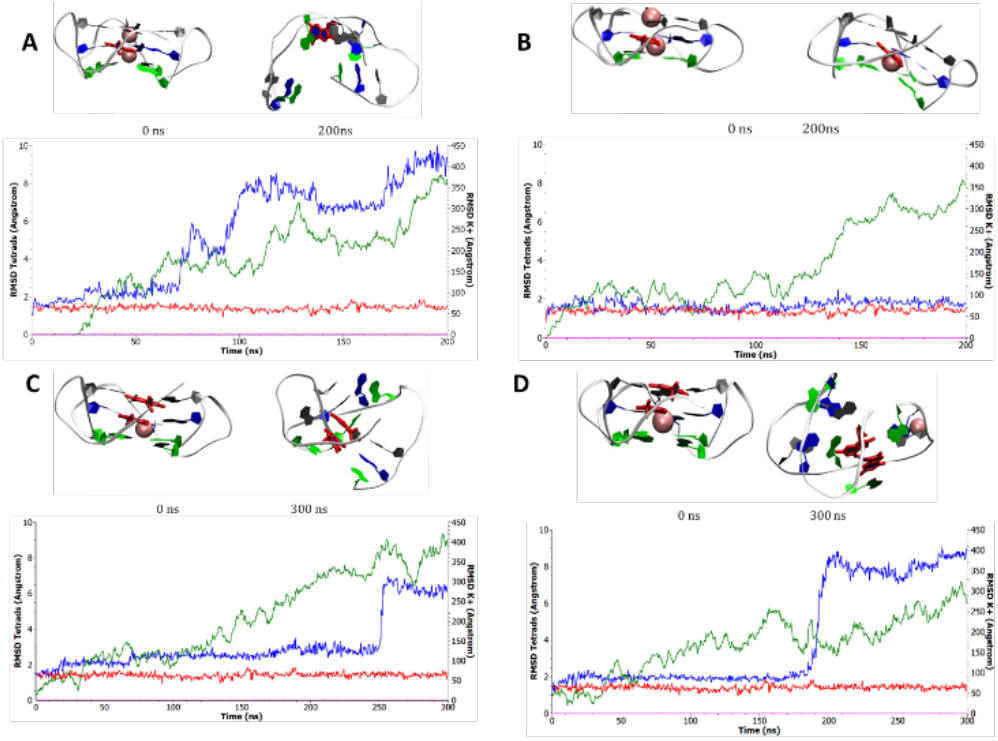
Starting and final conformation obtained for some representative G4s for simple (A) and (B) and double (C and D) lesions, 8-oxoG is placed at position 3 and 2-3, respectively leading either to conserved or disrupted G4s. The time evolution of the RMSD for the tetrads (green and blue curve) and for the central K^+^ ions (red and magenta) are also reported. Note the different scale for the two sets of curves.

**Table 1:** Summary of the main outcome of the MD simulations. (A) Globally conserved: the structure maintains the characteristic of a G4 persistently along the MD; (B) G4-like: only one tetrad is disrupted while the other maintain the G4 arrangement; (C) Disrupted: the G4 structure is totally lost.

|  | Number of simulation<br>(2 truns) | Number of K+ lost |  | Topology |  |  |
| --- | --- | --- | --- | --- | --- | --- |
|  |  | 1 | 2 | (A) Globaly conserved | (B) G4 like | (C) Disrupted |
| Parallel |  |  |  |  |  |  |
| 1 lesion | 8 | 0.88 | 12.50% | 75.00% | 18,75% | 6,25% |
| 2 lesion | 9 | 55.55% | 44.44% | 50.00% | 5.56% | 44.44% |
| Hybrid |  |  |  |  |  |  |
| 1 lesion | 2 | 0.00% | 0.00% | 100.00% | 100.00% | 0.00% |
| 2 lesion | 2 | 25.00% | 25.00% | 75.00% | 25.00% | 0.00% |

A crucial feature revealed by the MD simulations is the expulsion of one central K^+^ cation while preserving the folded quadruplex. In this case, our obtained structures look similar to the stable intermediate described by Zhang et al.^49^ the loss of the cation may be a consequence of the structural changes induced by the 8-oxoG lesion.

One of the most striking deformations observed concerns the arrangements of guanine O6 in the tetrads. Indeed, the introduction of one 8-oxoG changes the configuration of those atoms, which move from a nearly ideal square conformation to a rhomboidal arrangement, as illustrated in Figure 3, hence perturbing the tetrad arrangement.

**Figure 3:**
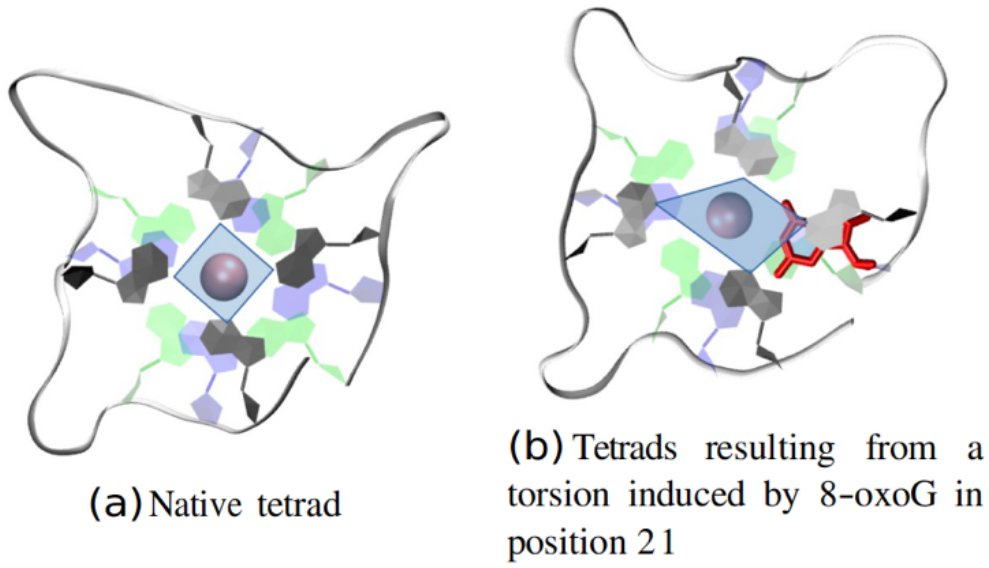
Deformation of the ideal disposition of the tetrad as an effect of the 8-oxoG lesion.

Unsurprisingly, and coherently with what observed for AP sites [add ref], the lesions on the central tetrad appears more disruptive. However, subtle sequence and position effects should be pointed out. As detailed in Figure 2, we can identify two peculiar situations: when 8-oxoG lesions is located on the A peripheral tetrad (position 2 and 14), we may still evidence the structural disturbance of the tetrad while the G4 structure is globally conserved. On the other hand, the lesion in position 16 induces a complete disturbance of the peripheral B tetrad. These observations cannot be explained solely as an effect of steric clash. Indeed, Giorgi et al.^50^ points out that an 8-oxoG strand is able to assemble in quartet forming other types of hydrogen bonds than those involving guanines. Hence, 8-oxoG is potentially capable of forming non-Hoogsteen hydrogen bonds with a neighboring nucleotide. This is confirmed by our simulations that clearly evidences the formation of persisting non-covalent interactions between 8-oxoG and the neighbouring guanines involving standard Hoogsteen hydrogen bonds (see SI). In addition, specific interactions can be pointed out involving either guanine and the cyclopenthenyl moiety of 8-oxoG in a fashion already pointed out by Giorgi et al.,^50^ and the interactions between H1 and H_2_1 of one nucleotide and the O6 atom of the second partner. Interestingly, this latter interaction, already evidenced experimentally by Bielskutè et al.,^44^ is formed not only between undamaged guanine and 8-oxoG, but also between two guanines or between two damaged nucleobases (see SI). While the specific outcome of the structural rearrangement strongly depends on the sequence and the position of the 8-oxoG lesions, it is clear that the tendency to maintain a G4 or G4-like structure is clearly emerging as the dominant motif when only one lesion is present, i.e. a situation that can be thought to be compatible with moderate oxidative stress conditions.

The sequence dependence observed in the case of a single 8-oxoG lesion is expected to emerge also in the case of a strand featuring two damages. However, as observed in the case of both double strand and G4 DNA in presence of cluster lesions, the coupling between the two damages may open further deformation paths resulting in different and more extreme structural outcomes. Indeed, as summarized in Table 1, in this case the majority of the MD trajectories leads to globally unfolded structures, that are also accompanied by the concomitant release of both central cations. Interestingly, while 3 trajectories still preserve the G4 arrangement, only in one case the G4-like conformation, with the unfolding of only one tetrad, with the release of only one cation, is observed. Differently from the case of one single lesion the possible different arrangements of the two cluster lesions grows combinatorically (Figure 1) including situations in which the damage occupy the same (horizontally arrangement) or different tetrads (vertical arrangement), giving rise to large differences in their coupling and hence in their effects. Unsurprisingly, it turns out that the most disruptive effect is found in the case of horizontally placed lesions, especially when they involve the central tetrad. Indeed, this situation results in the unfold of the first tetrad that is then accompanied with the destabilization of the central K^+^, their release and finally the unfolding. On the other hand, in the vertical arrangement one can observe a larger resistance to the lesion. This is also confirmed by the fact that for some of the conformations we obtained two different results for the two replicas, indicating a complex and rather rough free energy landscape that may lead to the coexistence of folded and unfolded structures. Finally, and interestingly, when the lesion occupies both peripheral tetrads, we observe the expulsion of a central cation, and a global reorganization which maintains two of the quartet in a G4-like conformation, made possible by the slight sliding of the remaining K^+^ to occupy the region between the two leaflets. This situation is also indicative of a general tendency implying that the 8-oxoG containing G4 seams resistant to the loss of one cation, and indeed folded structures in which only one of the cations is present are observed. Furthermore, the unfolding process, in almost all of the cases, is temporally initiated by the loss of a first cation, but necessitates the further expulsion of the second one to be completed. Of particular interest is the case reported in Figure 1D in which we observe two temporally well distinct events: the destabilization of the first tetrad that proceeds rather smoothly and a sharp transition leading to the unfolding of the second one. Snapshots extracted at important time of the trajectory are reported in Figure 4. Interestingly, the analysis of the trajectory points to a rather complex equilibrium with the bulk K^+^ ions that are initially stabilizing the tetrad even after the loss of the two central cations. However, this arrangement leads to a metastable state that rapidly collapses at around 190 ns due, once again to the interaction with the cations that forms a cluster around the G4s and leads to favorable electrostatic interactions with the electron-rich guanines. The cluster of K^+^ ions is still present during the first step of the unfolding as shown by the 204 ns conformation.

**Figure 4:**
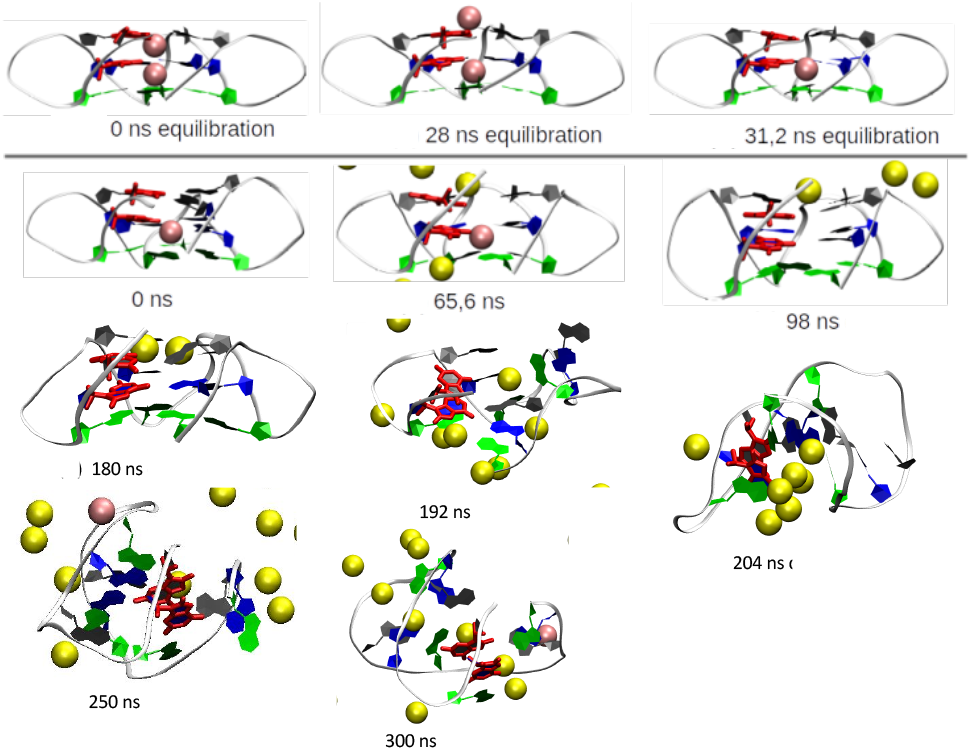
Snapshots representing the dynamic evolution of the double lesioned G4 reported in Figure 1D. Crystallographic K^+^, occupying the central canal are represented in pink, while the other cations are drawn yellow.

Thus, our results globally indicate a much larger structural destabilization produced by double-lesions on the G4 conformation, this can of course be easily justified by the much more extended structural perturbations induced by the coupled lesions. However, and compared to the presence of AP damages, 8-oxoG appears more innocent and even in the case of clustered lesions, i.e. in conditions of high or very high oxidative stress, the structural stability is higher, as witnessed by the presence of a non-negligible number of still folded sequence.

### Circular Dichroism

MD simulations are extremely advantageous in the study of the structural reorganizations induced as a consequence of DNA damages, also thanks to the molecular scale resolution that they can offer. Experimentally, a method of choice to unravel the complexity of even subtle structural modifications in biological ordered systems is electronic circular dichroism (ECD) spectroscopy.^19,51,52^ ECD spectra of the *h-Telo* G4 sequence, exposed to increasing concentrations of H_2_O_2_, in order to induce oxidative damage, were recorded. For the ECD studies, we have considered both a hybrid and a parallel folding, the former obtained in K^+^ rich buffered solution and the latter by including a crowding agent, such as poly-ethyleneglycol (PEG-200). The dichroism spectra are reported in Figure 5, while the corresponding UV absorption spectra can be found in SI.

**Figure 5.**
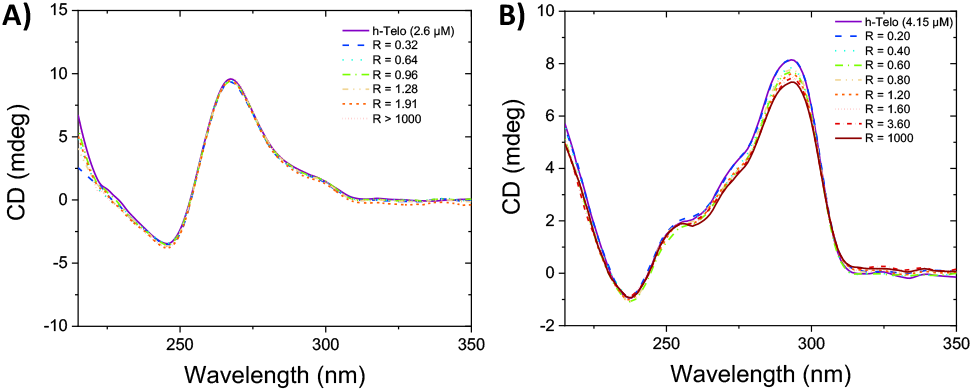
ECD spectra for parallel (A) and hybrid (B) *h-Telo* G4 DNA, recorded in the presence of increasing concentration of H_2_O_2_ (R = [H_2_O_2_]/[*h-Telo*]).

As can be seen in Figure 5, both arrangements show their typical spectroscopic signatures: a positive band near 265 nm and a negative peak centered around 240 nm for the parallel conformation and two positive peaks at 270 and 290 nm followed by a negative band near 240 nm for the hybrid strands. In both cases, the bands are globally maintained upon addition of H_2_O_2_. Indeed, while the ECD spectrum for the parallel arrangement is absolutely unchanged upon the addition of oxidant, only a very slight decrease of the intensity of the main positive band can be observed for the hybrid structure. Remarkably, all main spectroscopic features are maintained even in presence of a 1000-fold excess of H_2_O_2_. Due to the very strong sensitivity of ECD to secondary structure modifications in biological polymers, the results obtained can be safely interpreted in terms of a global conservation of the G4 structure and hence of their stability, coherently with the results obtained from the MD simulations. The minor decrease in intensity observed for the hybrid G4s can be attributed to the induction of some structural perturbation following single guanine to 8-oxoG oxidations that, as shown by the results of MD simulations, should be considered as minor. Of note, the slight perturbation of the ECD and of the UV absorption spectrum reported in SI can also be seen as an indirect confirmation of the formation of DNA lesions due to the effect of oxidation.

### In cellulo Immunofluorescence

Since the ECD spectra in solution confirm the stability of G4s exposed to oxidative stress, we went a step further in analyzing the behavior of healthy mammary epithelial cell lines, MCF10, exposed to H_2_O_2_. Via specific immunofluorescence assays, we identified and quantified both the presence of 8-oxoG and of G4s. Obviously, and differently from the solution case, in cellular media the amount of G4s after exposition to stress may be related to other factors in addition to the purely structural stability. These may include the influence of repair enzymes, the presence of complex signaling pathways and their cross-talk, and the global cellular response. Furthermore, a dependence upon the cellular cycle may also be observed and pinpointed. Hence, the immunofluorescence assays were also repeated in presence of antioxidants to assess for their effect. As displayed in Figure 6 and SI, we can observe that in absence of oxidative stress a minimal amount of 8-oxoG is present, on the other hand the addition of H_2_O_2_ leads to a very strong and statistically significant increase of its quantity. Unsurprisingly, the concomitant addition of H_2_O_2_ and antioxidants while still producing a significant amount of DNA lesions almost reduces its increase by half. As for the level of G4s in presence of antioxidants, H_2_O_2_ addition clearly leads to its increase, that is still statistically significant. However, differently from the 8-oxoG content, such increase appears as almost antioxidant independent. The increase of G4 content in condition of oxidative stress has already been documented in different cells lines and should be considered as a defense mechanism of the cell induced by the action of chaperone proteins.^53^ As such, it cannot be related uniquely to chemical and structural stability factors. However, the importance of the latter can be seen in the fact that the fluorescent labelling shows a rather large overlap for both 8-oxoG and G4s, indicating co-localization. Hence, we may safely conclude that while G4s can be seen as hotspots for DNA oxidative lesions, coherently with the high density of guanine, their structural stability helps in maintaining the global arrangement.

**Figure 6.**
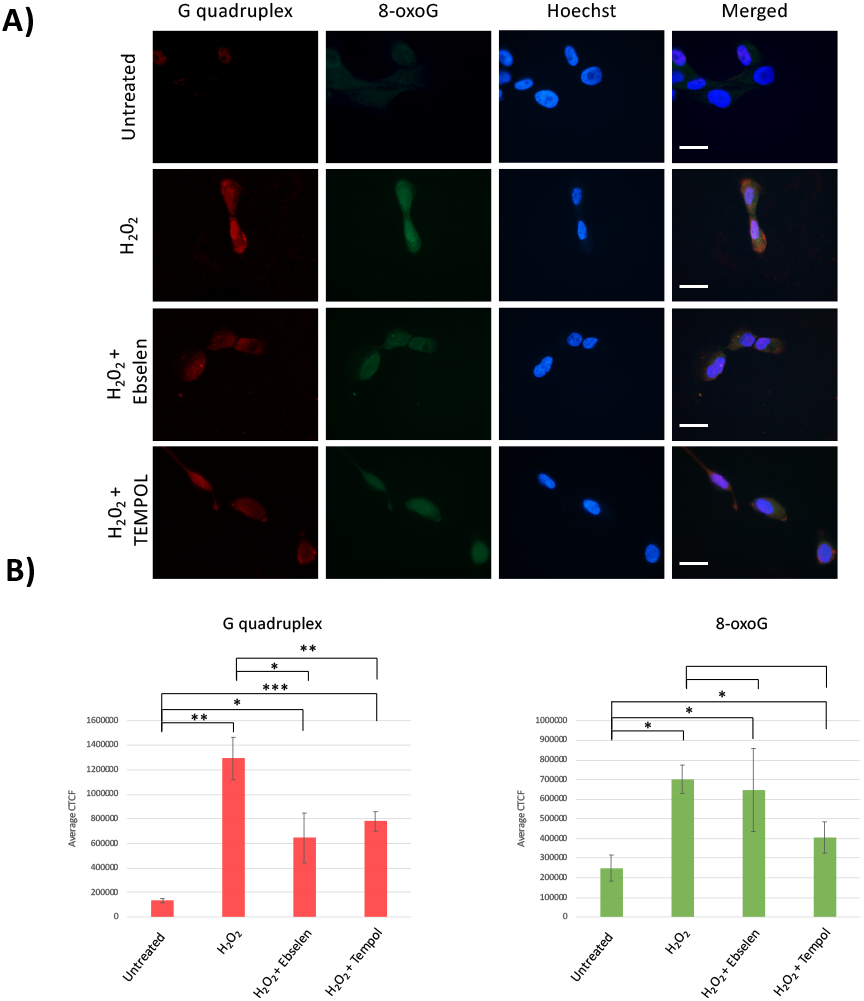
(A) Epifluorescence microscopy of MCF10a cells either untreated, treated with H_2_O_2_ or H_2_O_2_ and Ebselen or TEMPOL. Cells were fixed with paraformaldehyde and double stained with G quadruplex (red) and 8-oxoGuanine (green) antibodies. Scale bar represents 20 μm. (B) Quantification of fluorescence levels of immunofluorescence on MCF10a cells. Ten cells were counted per experiment. n = 3, T-test: * p<0,05, ** p<0,01.

## Conclusions

The combined use of our multiscale approach has allowed us to clearly sketch the scenario of the G4-DNA behavior in response to oxidative stress producing 8-oxoG. All results obtained, going from molecular modeling to cellular biology assays through spectroscopic studies, clearly point out a remarkable structural stability of the G4s to oxidative stress. MD simulations show that almost all the *h-Telo* sequences harboring an isolated damage are stable and do not undergo extended unfolding. Even the inclusion of a secondary lesion, while increasing the amount of unfolded sequences, is still characterized by a higher structurally stability compared to other lesions, such as AP sites. These results nicely support the interpretation of the virtually unchanged ECD spectra observed upon G4 titration and in the increase of the G4s amount in MCF10 cell lines treated with H_2_O_2_. Furthermore, the results of the cellular biology assays clearly show global nuclear localization of 8-oxoG and G4s, confirming that the chemical structural stability is a prerequisite for the cellular response to oxidative stress, i.e. the increase of the amount of G4s. These results allow us to sketch out some considerations on the biological role of G4-DNA. G4s are in fact known to act as regulators of the gene expression, with the most notable case of the *c-myc* oncogene, or as regulator of telomerase activity especially in the case of *h-Telo*. In particular, the inhibition of telomerase by DNA folding in G4 conformation helps to prevent cells to acquire immortality via the progressive shortening of telomeres. In conditions of strong oxidative stress, cells may be exposed to increased amount of DNA damages that may lead to mutations or even carcinogenesis. Hence, it is reasonable that the situation helping to confer immortality, and hence possible aggressivity and tumor-like phenotypes should be avoided. This can be indeed achieved by the stabilization of telomeric G4s. However, G4s being hot-spot for oxidative lesions due to the high guanine density, such a strategy undoubtedly requires a strong structural stability and resistance of the DNA conformation. From a chemical and biophysical point of view this is achieved by the fact that 8-oxoG, differently from AP sites, is still able to engage in non-Hoogsteen hydrogen-bonds with neighboring guanines, that while inducing a partial deformation of the tetrads maintain the global arrangement. This in turn is translated also in a much stronger propensity to the maintaining of the central cation, whose stabilizing role is crucial to keep the folded G4. Indeed, the case of AP lesions the global stability was achieved at the expense of one of the quartets, that was sacrificed to lead to a more extensive structural rearrangement. The higher stability is also witnessed by the fact that even the presence of clustered 8-oxoG, while obviously inducing a much stronger destabilization of the secondary structure, are in some resulting of preserved G4 or G4-like conformations.

With our work we have contributed to analyze the effects of oxidative lesions on the structural behavior of G4-DNA, clarifying also their biological role. In the future we plan to expand the present study considering the interactions of damaged G4s with protein partners, either BER repair enzymes or transcription factors. The structural effects of other lesions, such as strand breaks in the stability of the G4s arrangements would also be fully considered, since the latter can also be related to the effects of ionizing radiations.

## Experimental Section

### Molecular Dynamics Simulations

Each MD trajectory has been performed following the same protocol. In all the cases we have considered the *h-Telo* sequence folded in a parallel G4 arrangement as the starting point, to which 8-oxoG lesions have been manually added to specific positions. All the damaged DNA sequences have been solvated through a TIP3P box of water.^54^K^+^ are added to ensure the electroneutrality of the system; original K^+^ cations positioned inside G4 structure are conserved. A buffer of 10 Å of water is added to create the final octahedral box. Standards constants 300K and 1atm conditions are used to set up the dynamic simulations in the NPT ensemble. Amber ff99 force field including bsc1 corrections^55^ is used to describe DNA, while 8-oxoG potential is described by a specific force field designed by Bignon et al. in a previous work.^33^ Hydrogen mass repartitioning (HMR)^56^ is consistently applied to increase the non-water hydrogen mass, hence allowing the use of a 4 fs time step in combination with the Rattle and Shake algorithms.^57^ 1000 step of minimization are performed on the initial systems to remove bad contacts, followed by equilibration and thermalization for a total of 36 ns. All calculations were performed on two replica and continued until the RMSD of the designed G-quadruplex DNA is stable, i.e. between 200 ns and 300 ns. Trajectory calculations were performed using the NAMD software.^58^ All the MD trajectories have been analyzed and visualized using VMD.^59^

### Electronic Circular Dichroism

The *h-Telo* G4 sequence (5’-AGG GTT AGG GTT AGG GTT-3’) was purchased from IDT (Integrated DNA Technologies, Belgium) in HPLC purity grade. The oligonucleotide was dissolved in MilliQ water to yield a 100 μM stock solution. This was then diluted using 50 mM Tris-HCl/100 mM KCl buffer (pH 7.4) to the desired concentration. When needed, PEG-200 at 40% w/v was added to the buffer in order to obtain a parallel folding of the G4. The oligonucleotide was annealed heating the solutions up to 90 °C for 5 min and then by slowly cooling down to room temperature overnight. *h-Telo* concentration was checked measuring the absorbance at 260 nm and using 184000 L/(mol·cm) as extinction coefficient. Hydrogen peroxide concentration was determined by a redox titration with KMnO_4_. A stock solution of 0.982 M of H_2_O_2_ was kept in the fridge and used fresh for the *h-Telo* oxidation experiment. The ECD titrations were carried out by adding increasing amounts of properly diluted H_2_O_2_ to a solution of *h-Telo* at fixed concentration.

### Immunofluorescence Assays

Immunofluorescence assays were performed on MCF10a cells, a non-tumorigenic human breast epithelial cell line. MCF10a cells were cultured at 37°C, 5% CO_2_ in DMEM/F12, supplemented with 5% horse serum, 2 mM L-glutamine, 100 U/mL penicillin, 100 U/mL streptomycin, 10 μg/mL bovine insulin, 0,5 μg/mL hydrocortisone, 100 ng/mL cholera toxin and 20 ng/mL hEGF. MCF10a cells were cultured on a glass slide in complete medium for 24h before treatment. Cells were treated with either 200 μM H_2_O_2_ (H1009, Sigma-Aldrich) for 1h with or without antioxidant, 50 μM Ebselen (E3520, Sigma Aldrich) or 3 mM TEMPOL (176141, Sigma Aldrich). Cells were fixed in 4% paraformaldehyde over 20 minutes at room temperature, then blocked and permeabilized with PBS-containing 2% BSA/0.2% Triton X-100. Cells were incubated with anti-DNA/RNA G-quadruplex [BG4] primary antibody (Ab00174-1.1, Absolute Antibody) diluted at 1:200 in PBS-containing 2% BSA for 30 minutes at 37°C. After three washes in PBS, the cells were incubated with Alexa Fluor 594– conjugated goat anti mouse secondary antibody (A-11032, Invitrogen), diluted at 1:500 in PBS-containing 2% BSA for 20 minutes at 37°C. The cells were washed three times in the washing buffer 1X of OxyDNA Assay Kit (500095, Calbiochem) and incubated overnight at 37°C in FITC-conjugated anti 8-oxoguanine (500095, Calbiochem) diluted at 1:100 in washing buffer 1X. After three washes with washing buffer, nuclei were stained with Hoechst diluted at 1:10,000 in PBS. The cells were then mounted in antifading medium (FluorSafe; Merck) and observed with an epifluorescence microscope Eclipse 80i with ×100 oil immersion objective (Nikon). Images were collected with a digital camera (Nikon, DS-Ri1) with the same exposure time for all the conditions. Cells fluorescence levels have been assessed following measuring cell fluorescence using ImageJ entry in The Open Lab Book, contributed by Luke Hammond, QBI, The University of Queensland, Australia Hammond 48. Cells were selected on ImageJ using the drawing tool, and their area, integrated density and mean grey value were measured. Regions around the cell were selected and measured to determine the background. Corrected total cell fluorescence (CTCF) was determined as the integrated density minus the product between the area of the selected cell and the mean fluorescence of the background reading. Graphs were made using mean CTCF value for each condition, error bars correspond to SEM.

## Acknowledgements

Support from the Ministero dell’Università e Ricerca Scientifica e Tecnologica and Università di Palermo, Italy, and Université de Lorraine and CNRS, France, are gratefully acknowledged. Tom Miclot thanks University of Palermo for funding a joint Ph.D. program. CINECA - SCAI national computing center is acknowledged for graciously providing access to computational resource. Part of the calculations have been performed on the local LPCT computational resources and on the regional ExpLor mesocenter in the framework of the project “Dancing Under the Light”.

## Entry for the Table of Contents

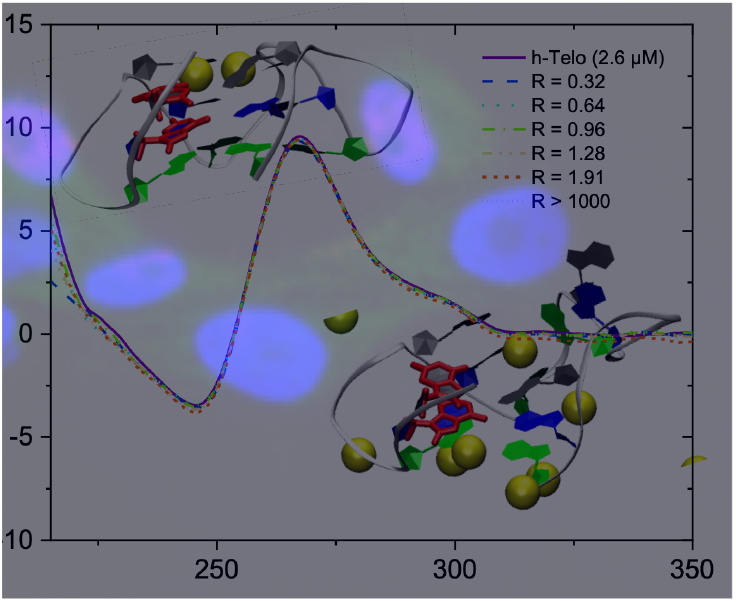

The structural stability of Guanine quadruplexes in presence of oxidative lesions is revealed using molecular simulation, CD spectroscopy and immunofluorescence.

Institute and/or researcher Twitter usernames: @AntonioMonari, @baronegiampaolo, @Al3XI0

## Notes

### Competing Interest Statement

The authors have declared no competing interest.

