## Supplementary Information for "Forever Young: Structural Stability of Telomeric Guanine-Quadruplexes in Presence of Oxidative DNA Lesions"

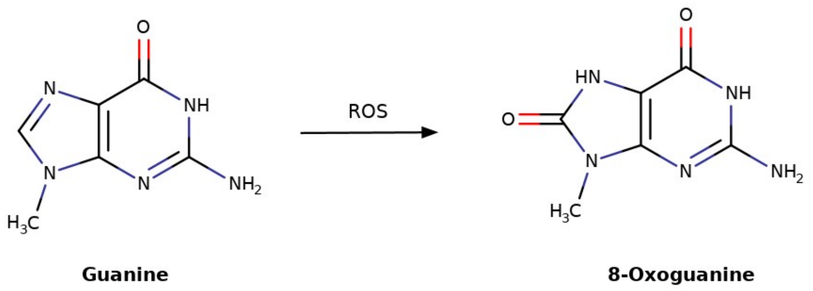


Figure S1: Molecular sketch of guanine and 8-oxoG.

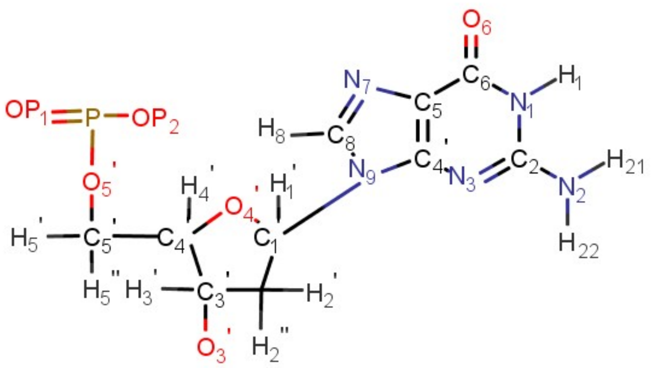

Figure S2: Naming scheme for the atoms in guanine coherent with Amber force field.

### Interactions between 8-oxoG and Guanine

Figure S3: Main classes of H-bond interactions found during the MD simulation. The 2D-chemical formula, naming conventions, and time series are provided.

<table>
<tr>
<td>(a) Standard Hoogsten H-bonds: G-G<br>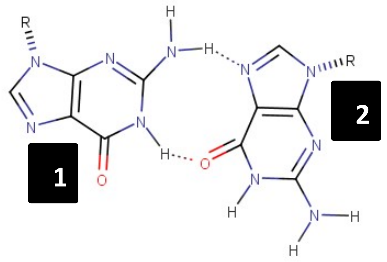
</td>
<td>- H1 of the guanine 1 and O6 of the guanine 2.<br>- H21 of the guanie 1 and N7 of the guanine 2.</td>
<td>///</td>
</tr>
<tr>
<td>(b) Standard G4 bonds: G-8oxoG<br>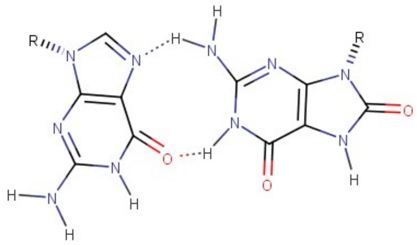</td>
<td>The standard G4 interactions can be established between a guanine and an 8-oxoG involving:<br>- H1 of the 8-oxoG and O6 of the guanine.<br>- H 21 of the 8-oxoG and N7 of the guanine.</td>
<td>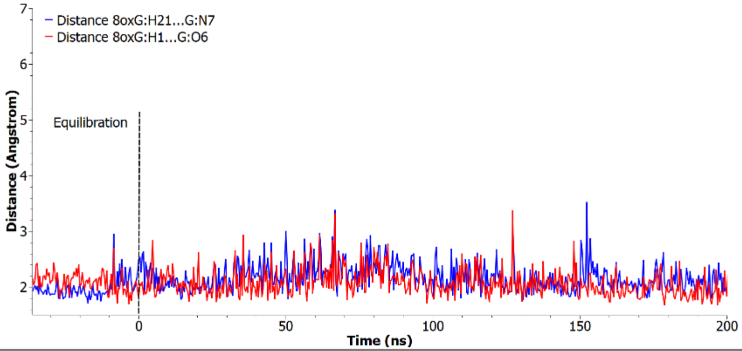
</td>
</tr>
<tr>
<td>(c) Standard 8-oxoG tetrad bonds : G-8oxoG<br>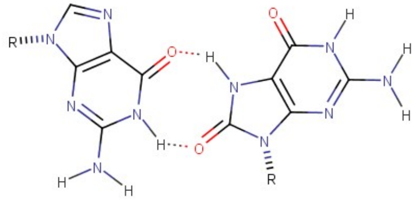</td>
<td>8-oxoG is able to form tetrads by interacting with the cyclopenthenyl moiety of guanine involving hydrogen-bonds between:<br>- H7 of the 8-oxoG and O6 of the guanine.<br>- O8 of the 8-oxoG and H1 of the guanine.</td>
<td>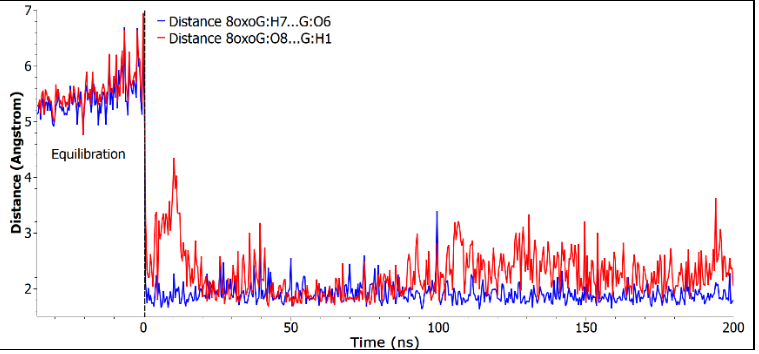
</td>
</tr>
</table>

<table>
<tr>
<td>(d) Non standard G4 bonds<br>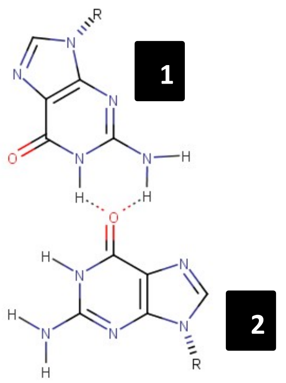</td>
<td>Two guanines are bonded with non standard interactions involving:<br>- H1/H21 of the guanine 1 and O6 of the guanine 2.<br>It is possible that this interaction is less strong than those found in the standard G4 bond.</td>
<td>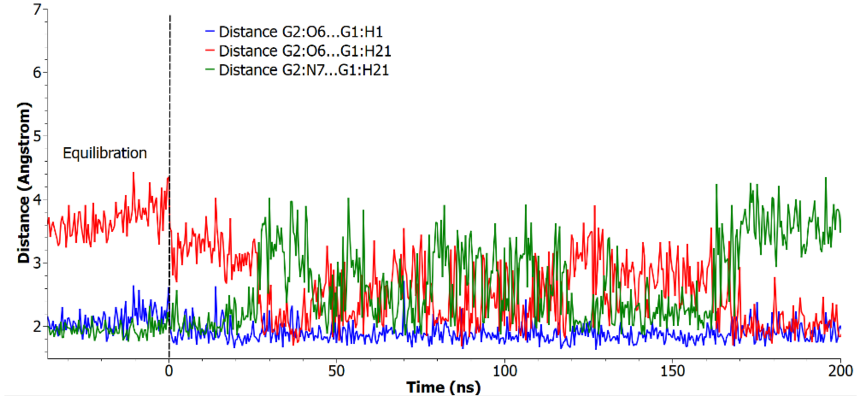</td>
</tr>
<tr>
<td>(e) Bond type :<br>G:H1/H21-8oxoG:O6<br>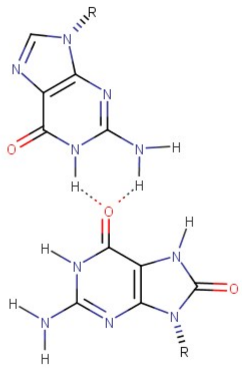</td>
<td>It establishes hydrogen interactions between :<br>- H1/H21 of the guanine and O6 of the 8-oxoG.<br>Structuraly, there is a steric clash between G:H21 and 8oxoG:H7. This explain why there is a shift between the two nucleotides leading to the inteaction bewteen G:H21 and 8oxoG:O6.</td>
<td>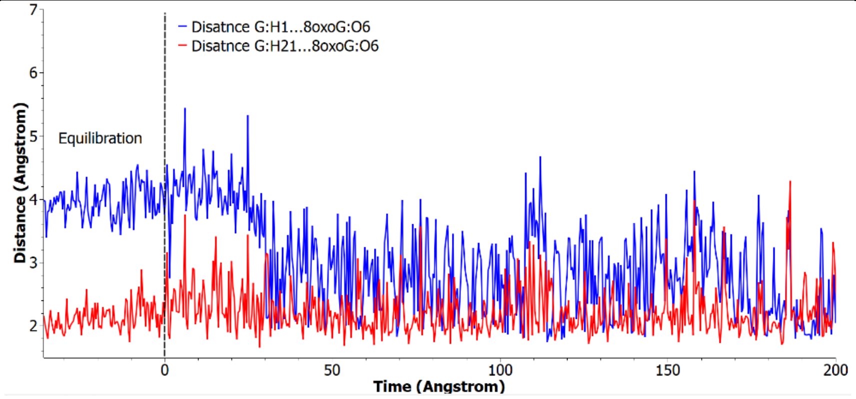</td>
</tr>
</table>

| | | |
|---|---|---|
| (f) Bond type : 8oxoG:H1/H21-8oxoG:O6 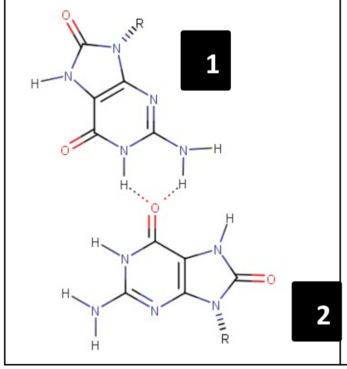  | Two 8-oxoG are bonded with non standard interactions. It establishes hydrogen interactions between :<br>- H1/H21 of the 8-oxoG 1 and O6 of the 8-oxoG 2. | 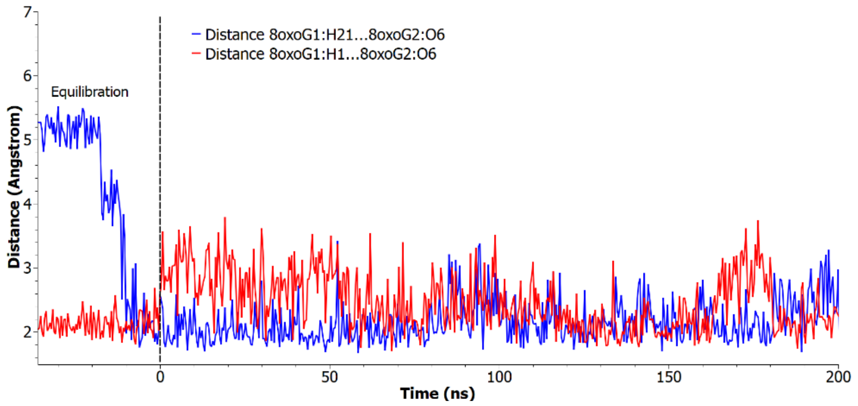  |

### Molecular dynamic simulations - Parallel G4 - Single lesion

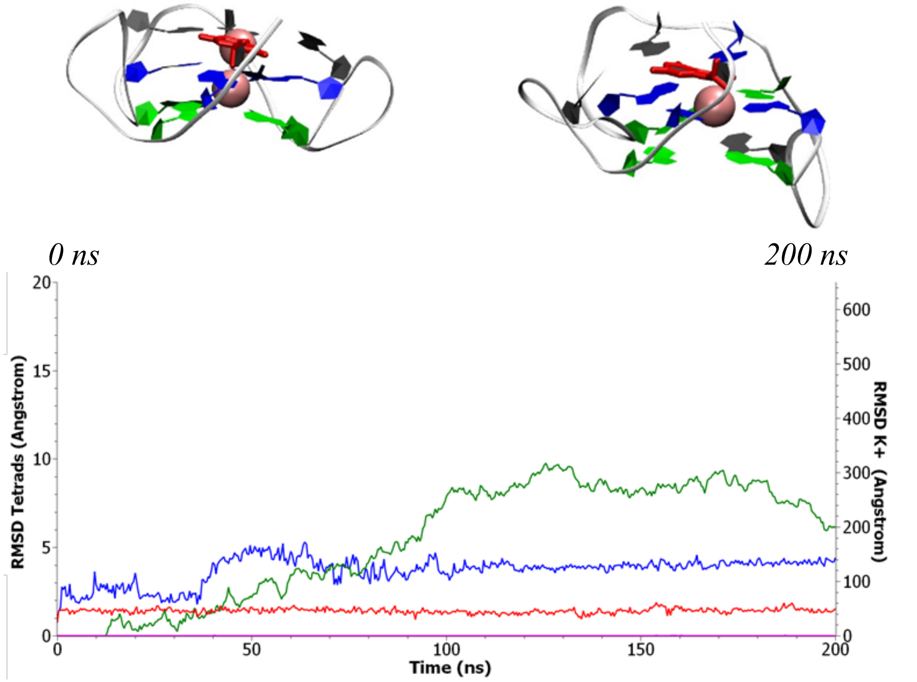


Run 1

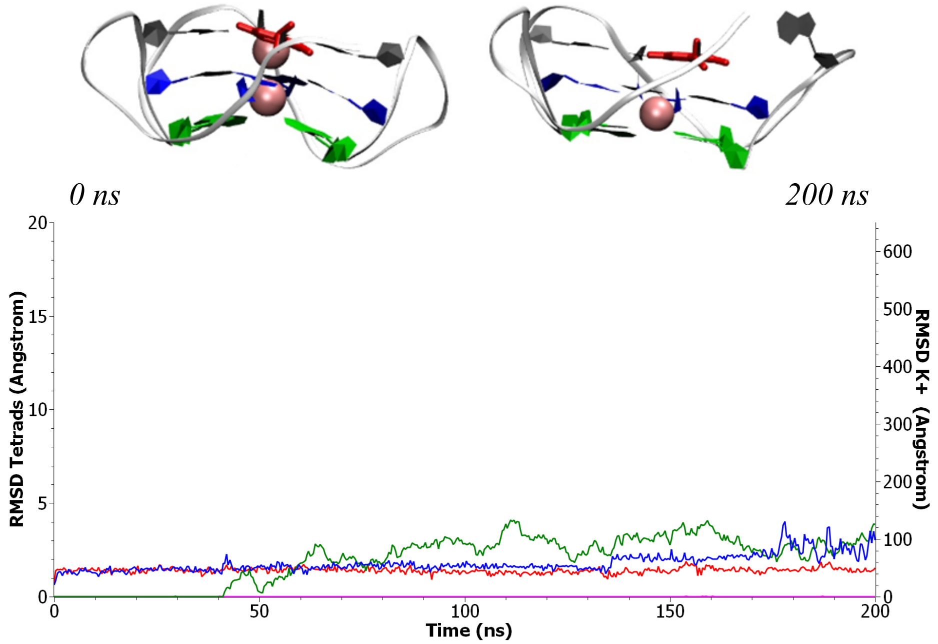


Run 2

Figure S4: Representative snapshots and time series of the RMSD for the two simulations involving parallel G4 presenting a lesion at position 2. *Red: RMSD of tetrads for the full undamaged structure. Blue: RMSD of tetrads for the full damaged structure. Pink: RMSD of $K^+$ for undamaged structure. Green: RMSD of $K^+$ for the damaged structure.*

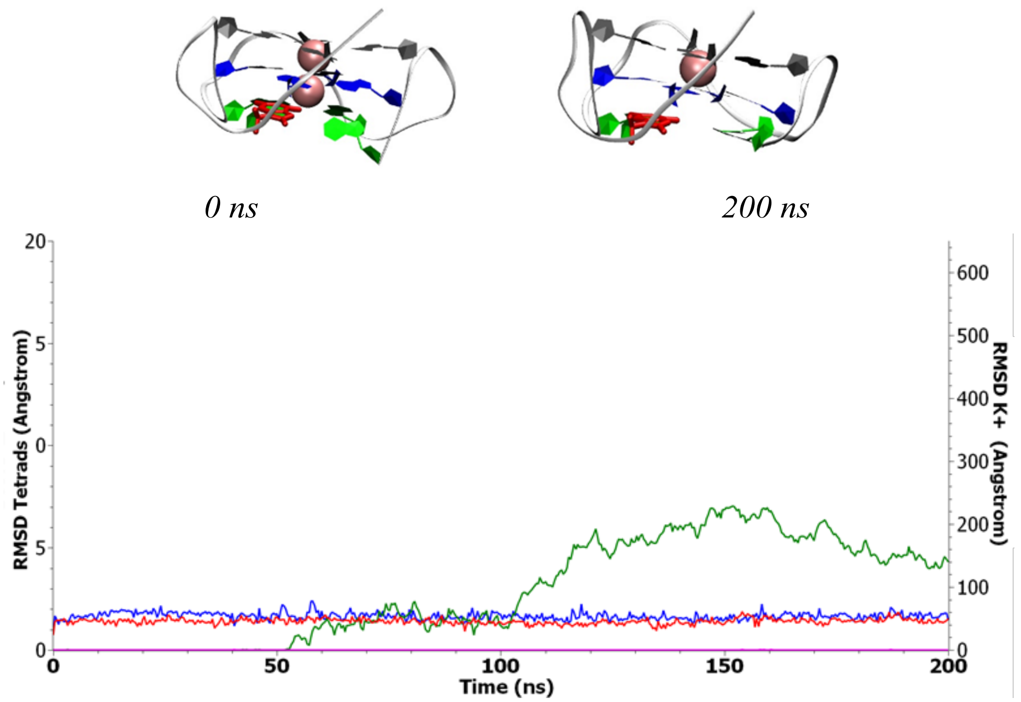


Run 1

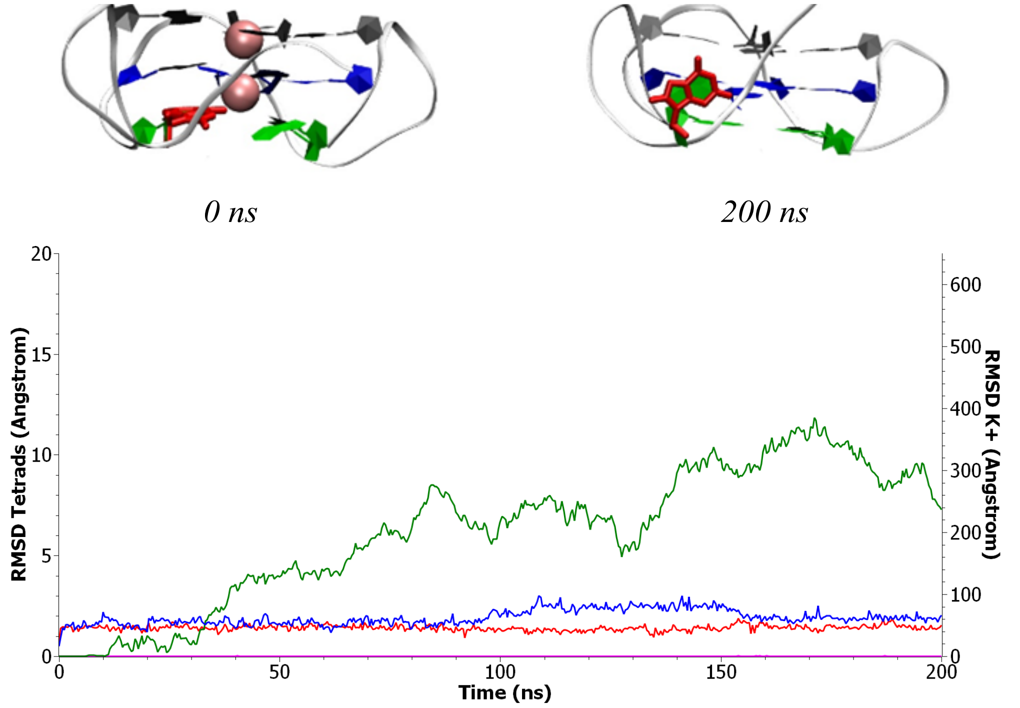


Run 2

Figure S5: Representative snapshots and time series of the RMSD for the two simulations involving parallel G4 presenting a lesion at position 4. *Red: RMSD of tetrads for the full undamaged structure. Blue: RMSD of tetrads for the full damaged structure. Pink: RMSD of $K^+$ for undamaged structure. Green: RMSD of $K^+$ for the damaged structure.*

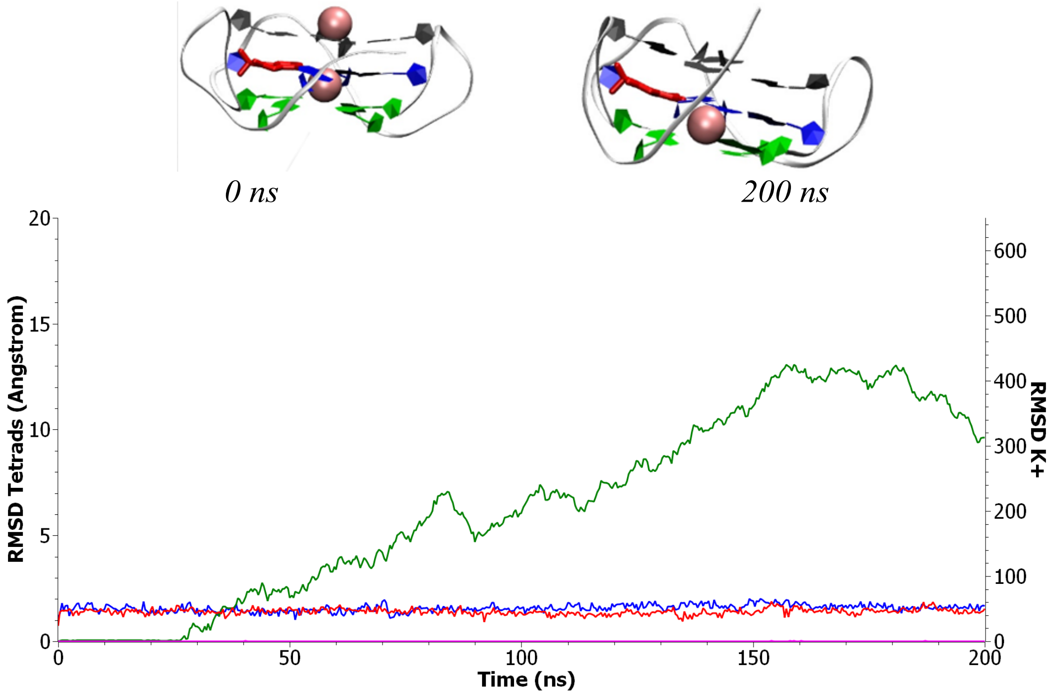


Run 1

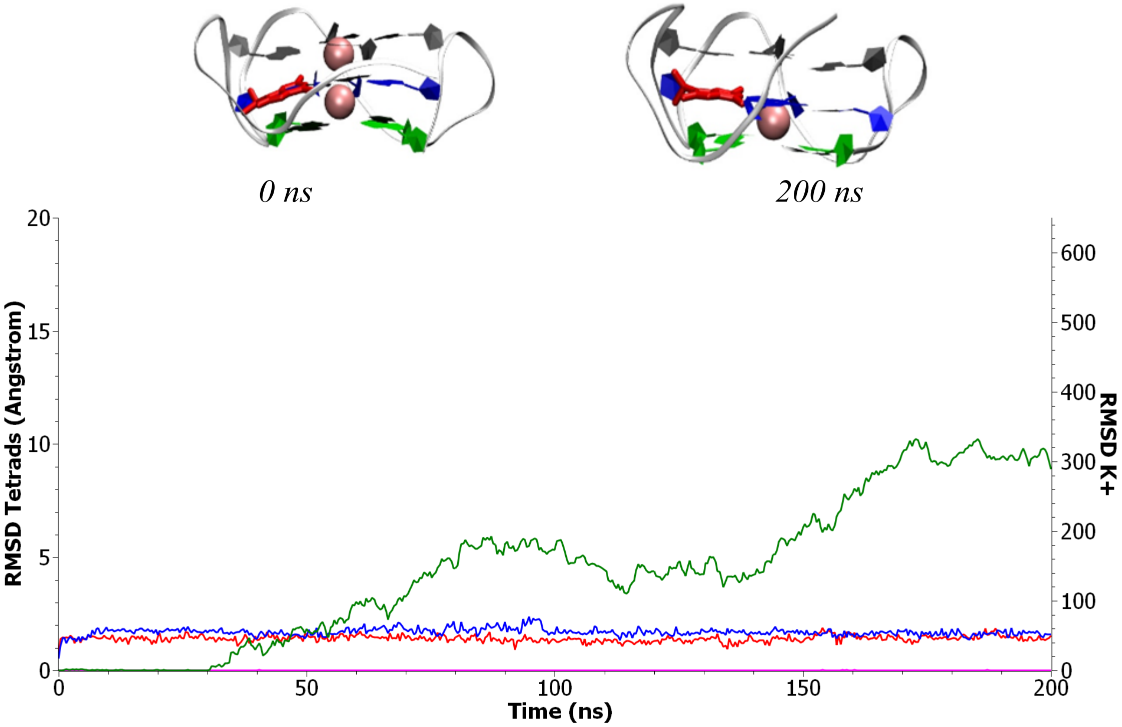


Run 2

Figure S6: Representative snapshots and time series of the RMSD for the two simulations involving parallel G4 presenting a lesion at position 9. *Red: RMSD of tetrads for the full undamaged structure. Blue: RMSD of tetrads for the full damaged structure. Pink: RMSD of $K^+$ for undamaged structure. Green: RMSD of $K^+$ for the damaged structure.*

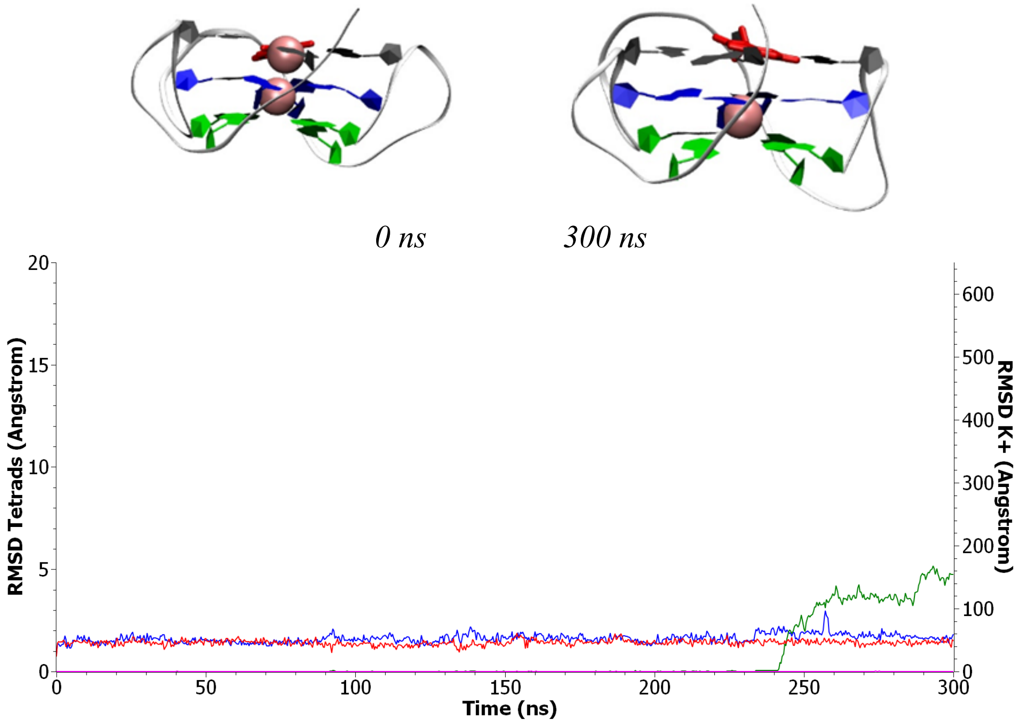


Run 1

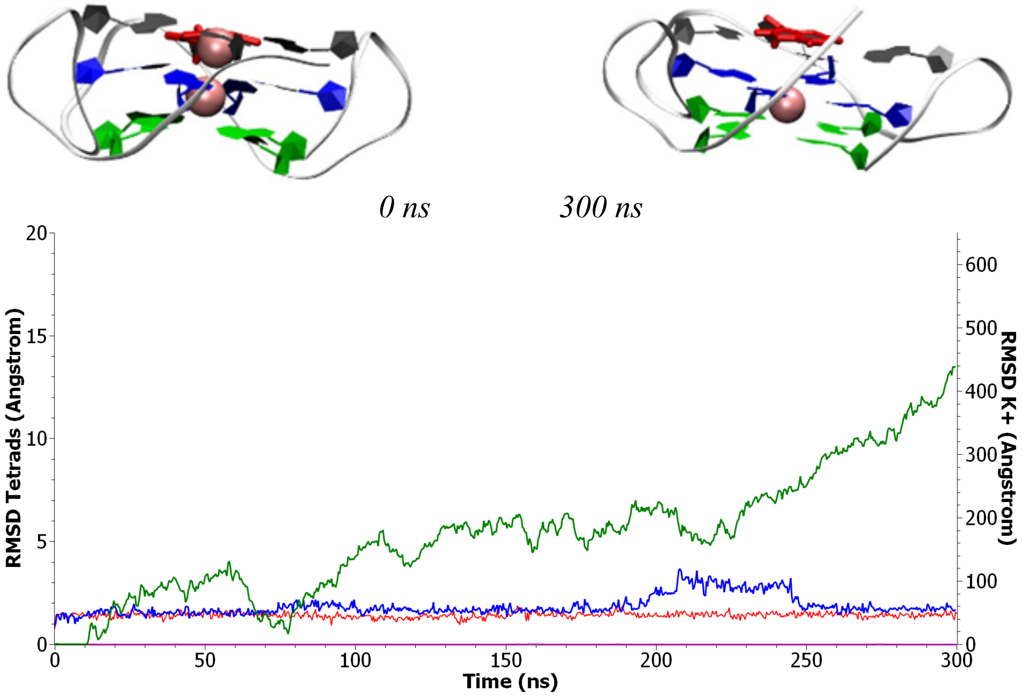


Run 2

Figure S7: Representative snapshots and time series of the RMSD for the two simulations involving parallel G4 presenting a lesion at position 14. *Red: RMSD of tetrads for the full undamaged structure. Blue: RMSD of tetrads for the full damaged structure. Pink: RMSD of $K^+$ for undamaged structure. Green: RMSD of $K^+$ for the damaged structure.*

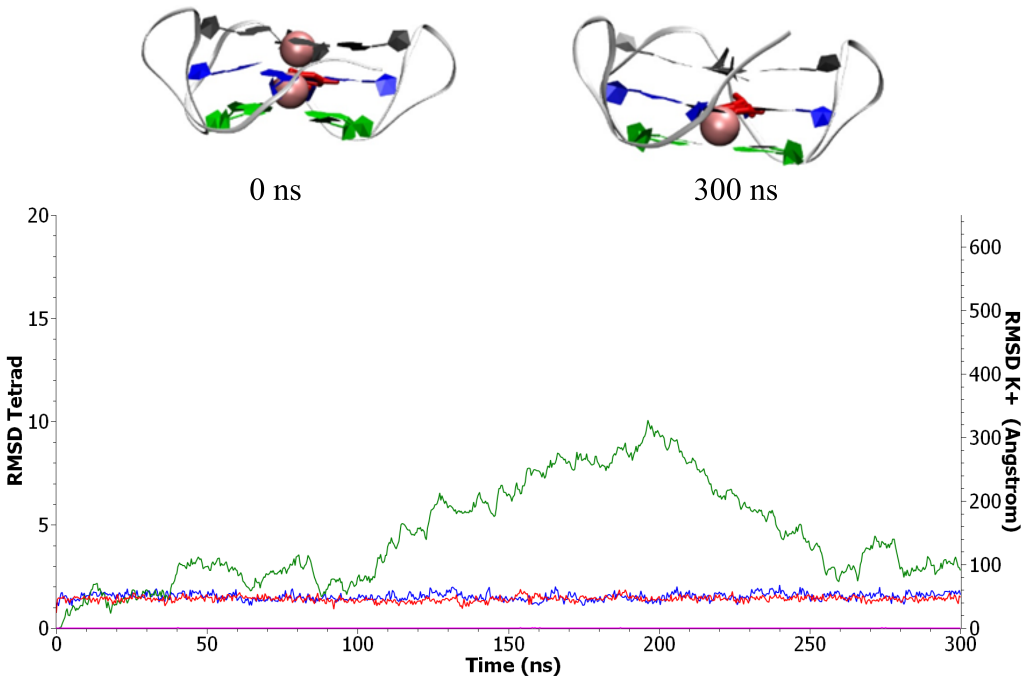


Run 1

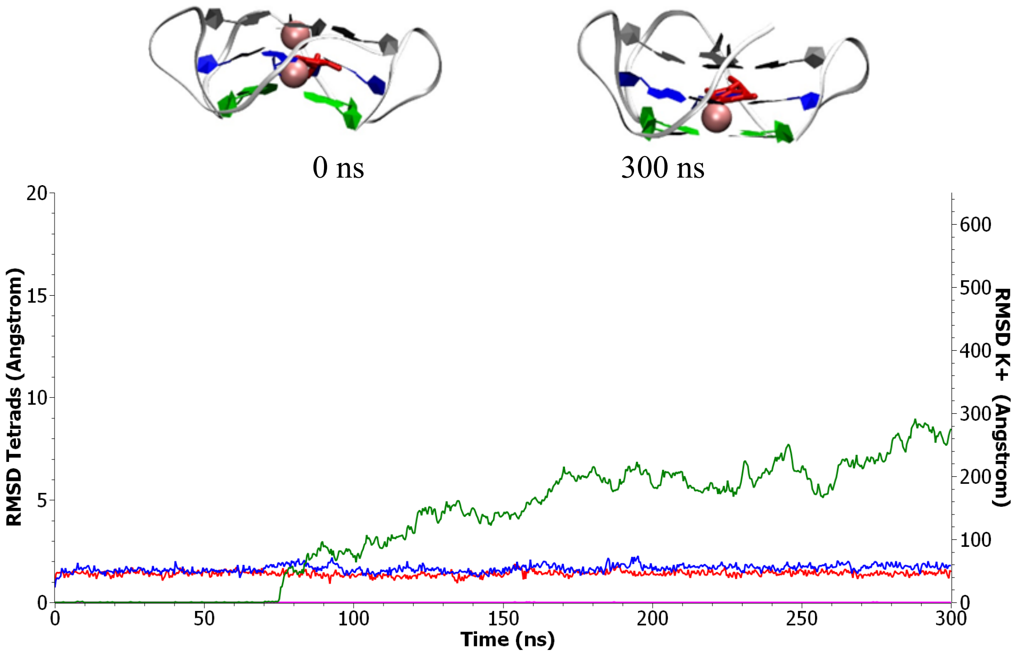


Run 2

Figure S8: Representative snapshots and time series of the RMSD for the two simulations involving parallel G4 presenting a lesion at position 15. *Red: RMSD of tetrads for the full undamaged structure. Blue: RMSD of tetrads for the full damaged structure. Pink: RMSD of $K^+$ for undamaged structure. Green: RMSD of $K^+$ for the damaged structure.*

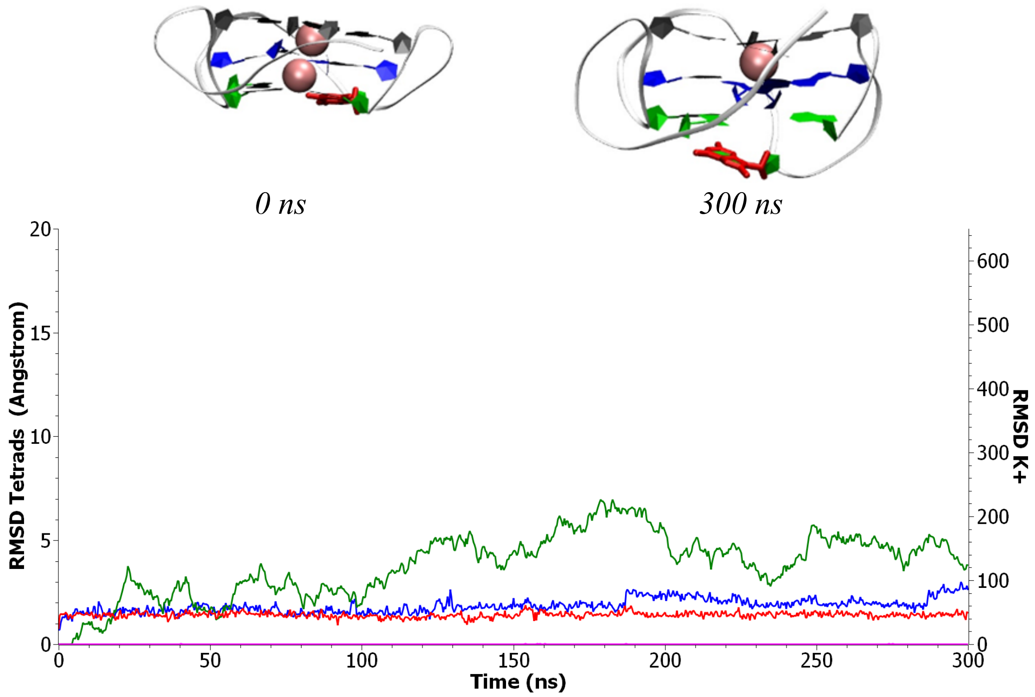


Run 1

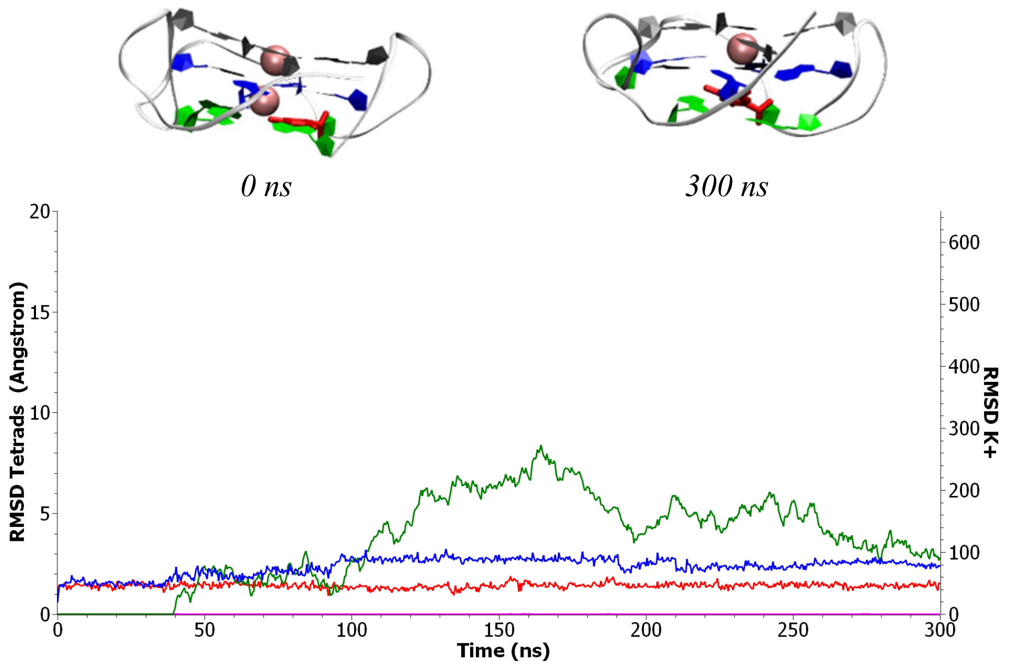


Run 2

Figure S9: Representative snapshots and time series of the RMSD for the two simulations involving parallel G4 presenting a lesion at position 16. *Red: RMSD of tetrads for the full undamaged structure. Blue: RMSD of tetrads for the full damaged structure. Pink: RMSD of $K^+$ for undamaged structure. Green: RMSD of $K^+$ for the damaged structure.*

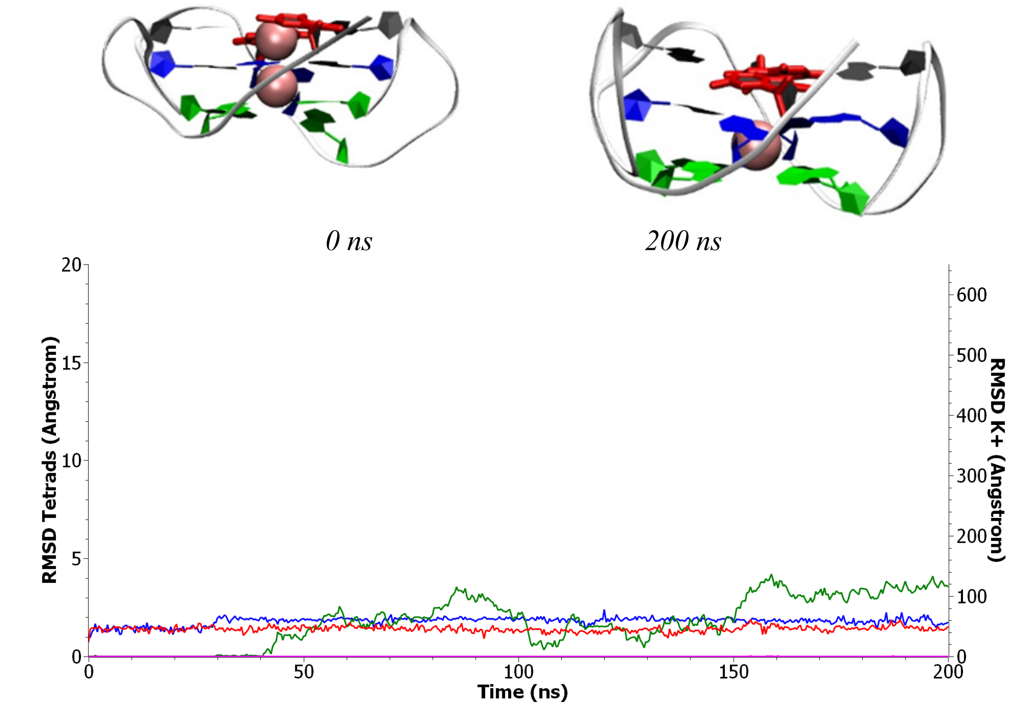


Run 1

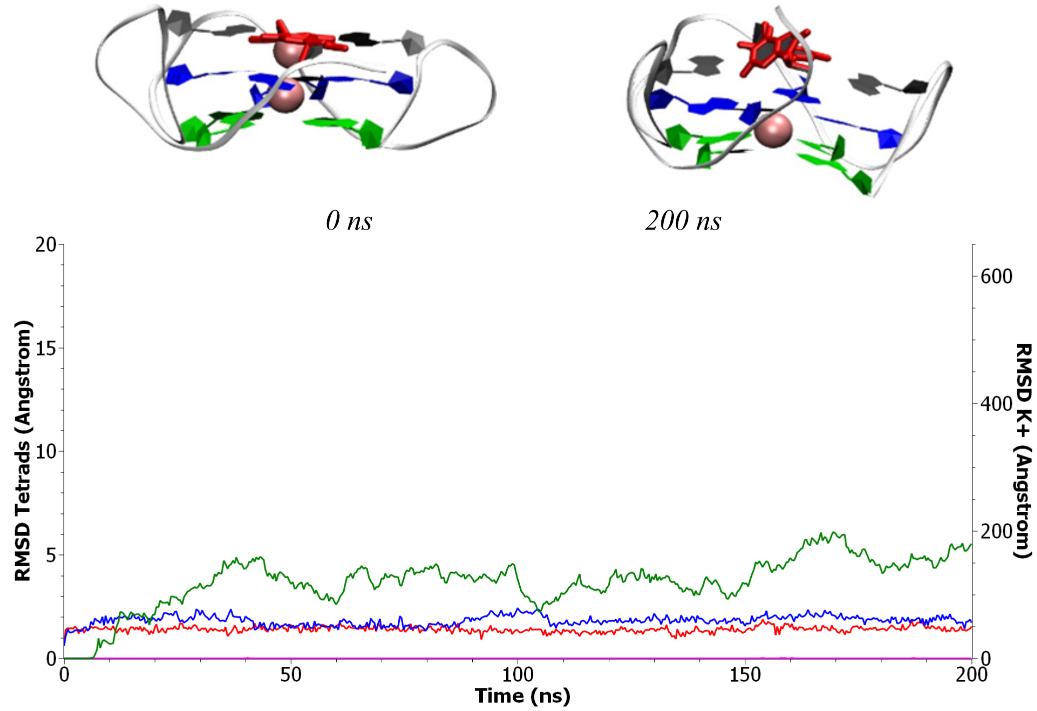


Run 2

Figure S10: Representative snapshots and time series of the RMSD for the two simulations involving parallel G4 presenting a lesion at position 21. *Red: RMSD of tetrads for the full undamaged structure. Blue: RMSD of tetrads for the full damaged structure. Pink: RMSD of $K^+$ for undamaged structure. Green: RMSD of $K^+$ for the damaged structure.*

### Molecular dynamic simulations - Parallel G4 - Double lesions

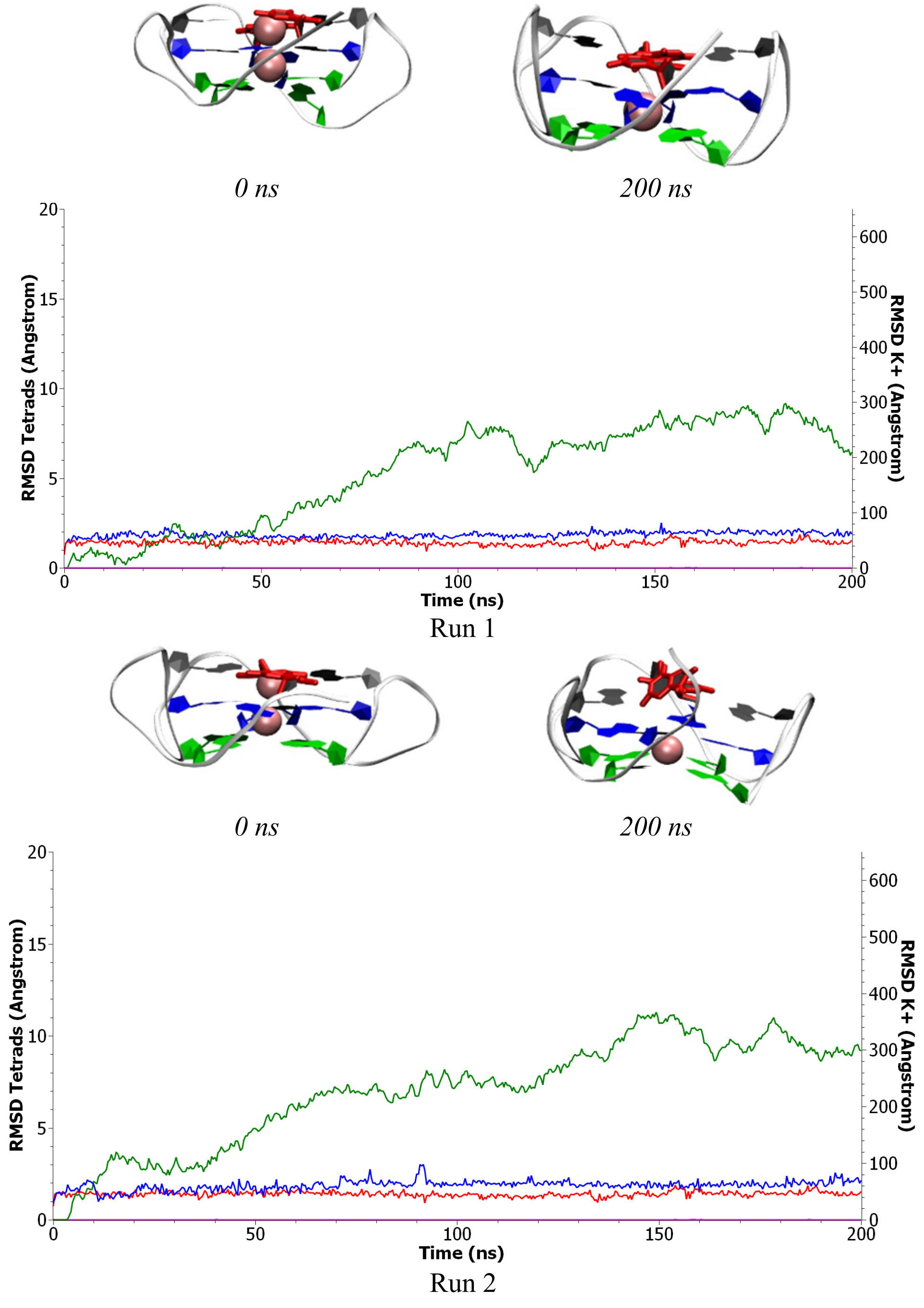


Figure S11: Representative snapshots and time series of the RMSD for the two simulations involving parallel G4 presenting double lesions at position 2 and 14. *Red: RMSD of tetrads for the full undamaged structure. Blue: RMSD of tetrads for the full damaged structure. Pink: RMSD of $K^+$ for undamaged structure. Green: RMSD of $K^+$ for the damaged structure.*

.

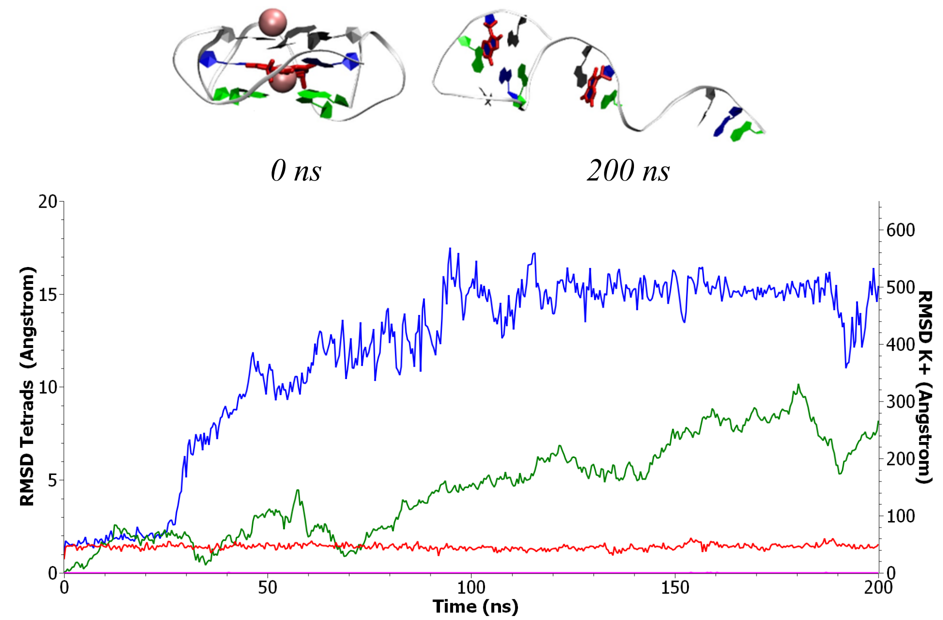


Run 1

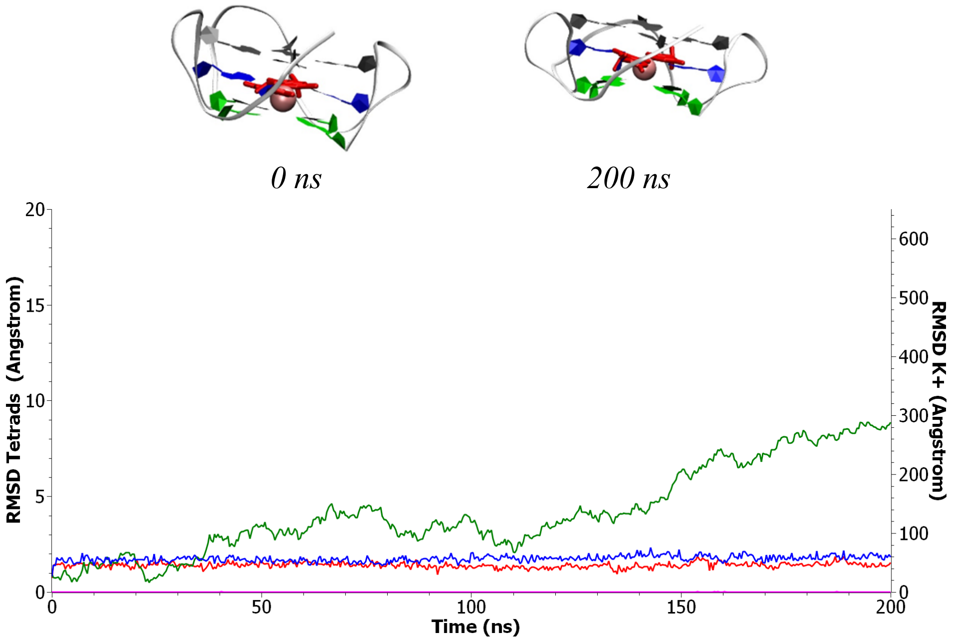


Run 2

Figure S12: Representative snapshots and time series of the RMSD for the two simulations involving parallel G4 presenting double lesions at position 3 and 15. *Red: RMSD of tetrads for the full undamaged structure. Blue: RMSD of tetrads for the full damaged structure. Pink: RMSD of* $K^+$ *for undamaged structure. Green: RMSD of* $K^+$ *for the damaged structure.*

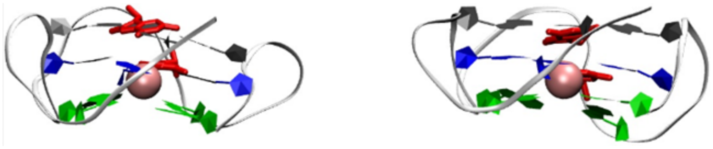

*0 ns* *300ns*

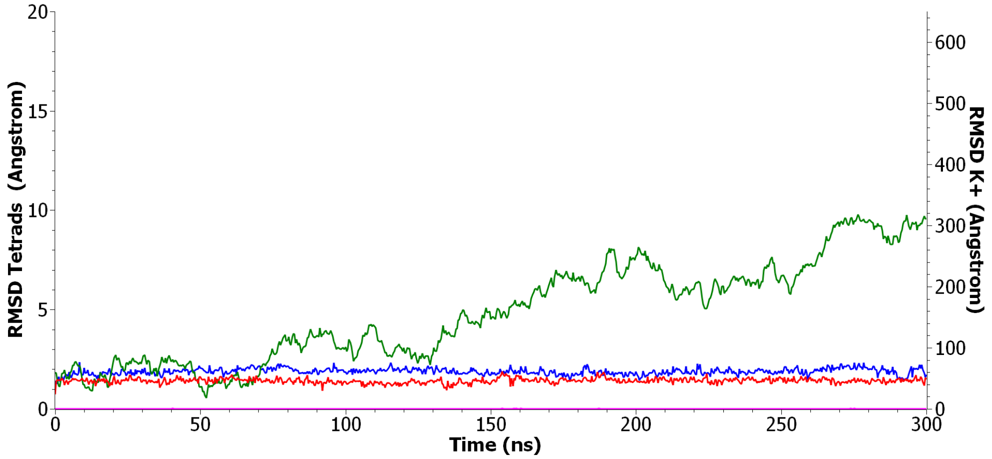


Run 1

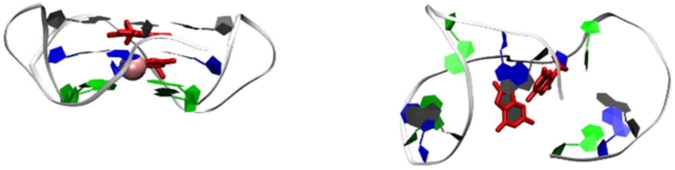

*0 ns* *300ns*

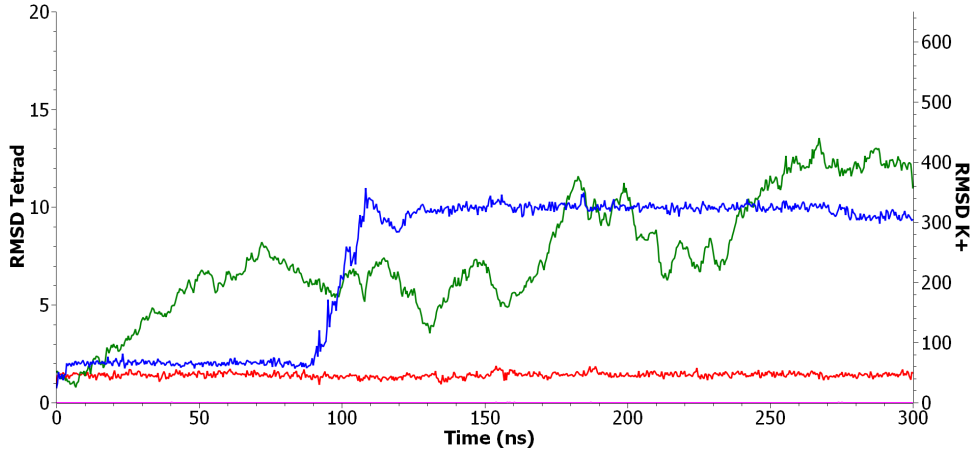


Run 2

Figure S13: Representative snapshots and time series of the RMSD for the two simulations involving parallel G4 presenting double lesions at position 14 and 15. *Red: RMSD of tetrads for the full undamaged structure. Blue: RMSD of tetrads for the full damaged structure. Pink: RMSD of $K^+$ for undamaged structure. Green: RMSD of $K^+$ for the damaged structure.*

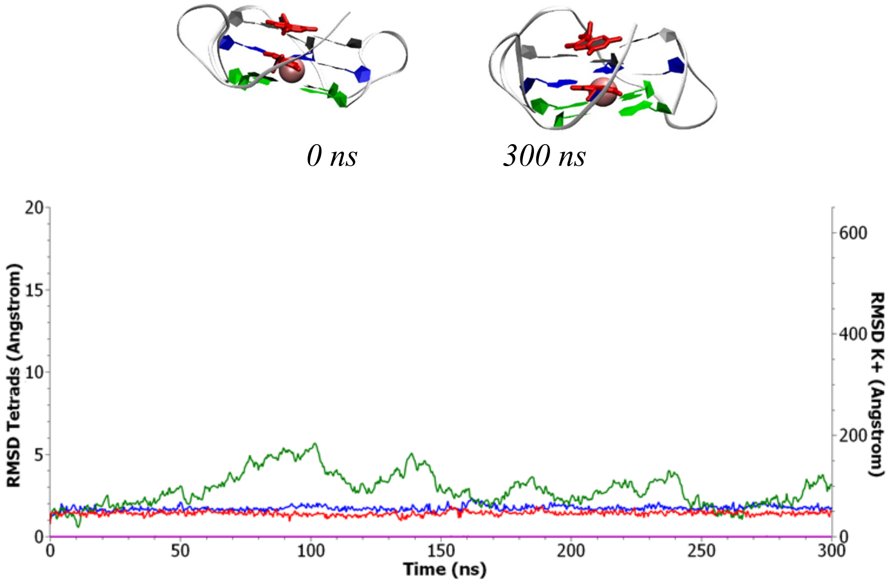


Run 1

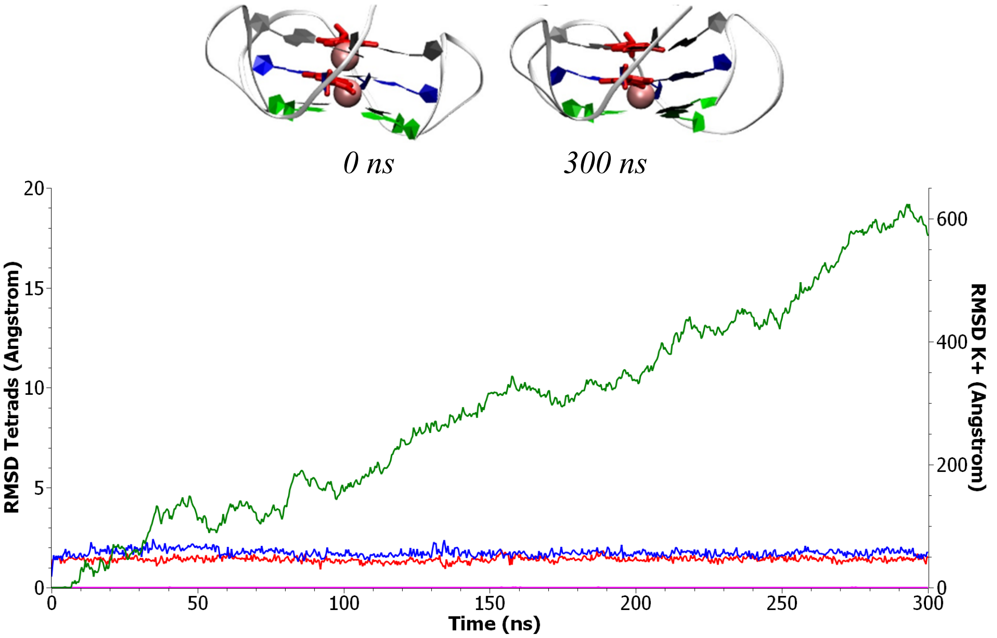


Run 2

Figure S14: Representative snapshots and time series of the RMSD for the two simulations involving parallel G4 presenting double lesions at position 3 and 14. *Red: RMSD of tetrads for the full undamaged structure. Blue: RMSD of tetrads for the full damaged structure. Pink: RMSD of $K^+$ for undamaged structure. Green: RMSD of $K^+$ for the damaged structure.*

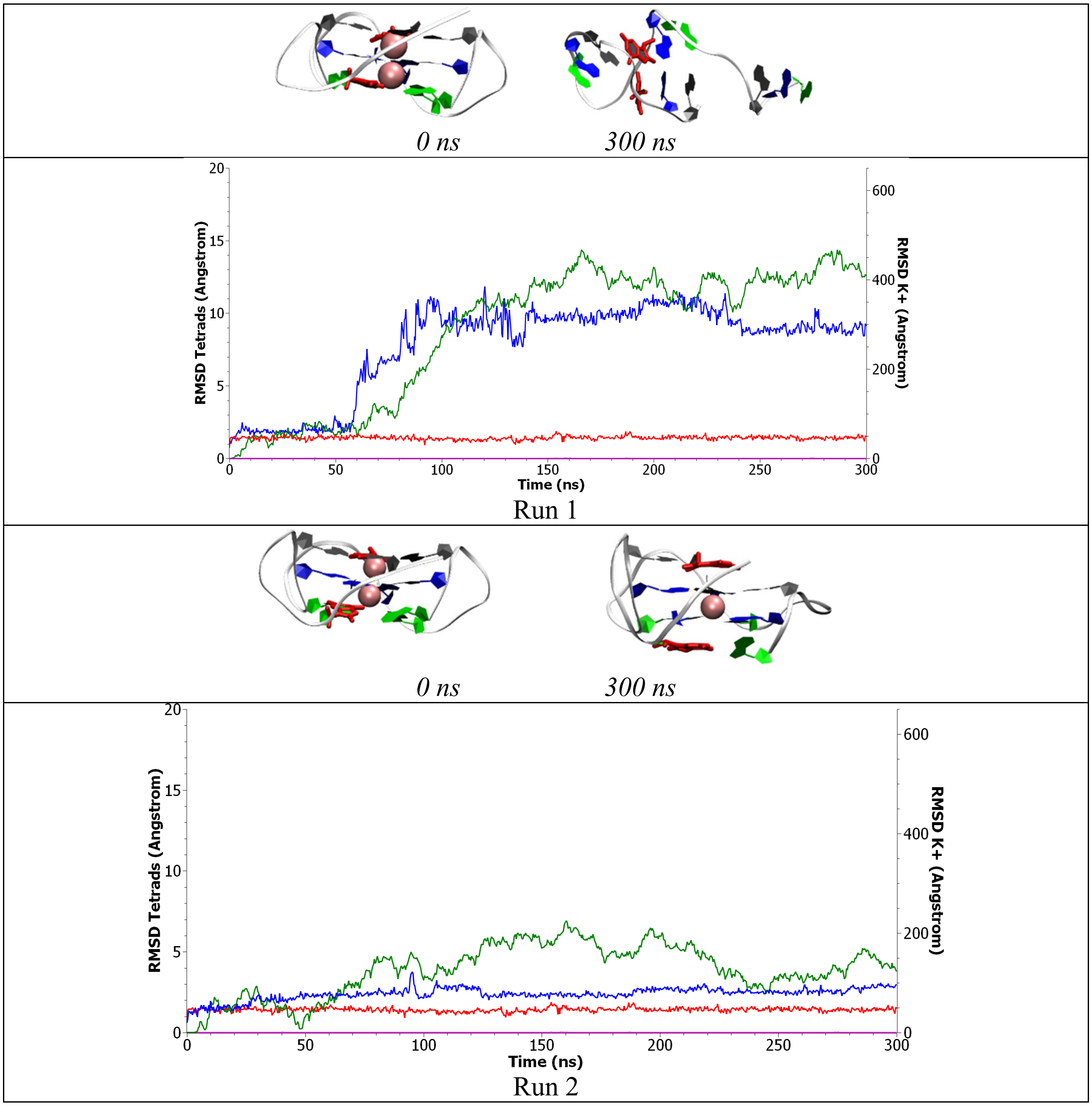


Figure S15: Representative snapshots and time series of the RMSD for the two simulations involving parallel G4 presenting double lesions at position 4 and 14. *Red: RMSD of tetrads for the full undamaged structure. Blue: RMSD of tetrads for the full damaged structure. Pink: RMSD of $K^+$ for undamaged structure. Green: RMSD of $K^+$ for the damaged structure.*

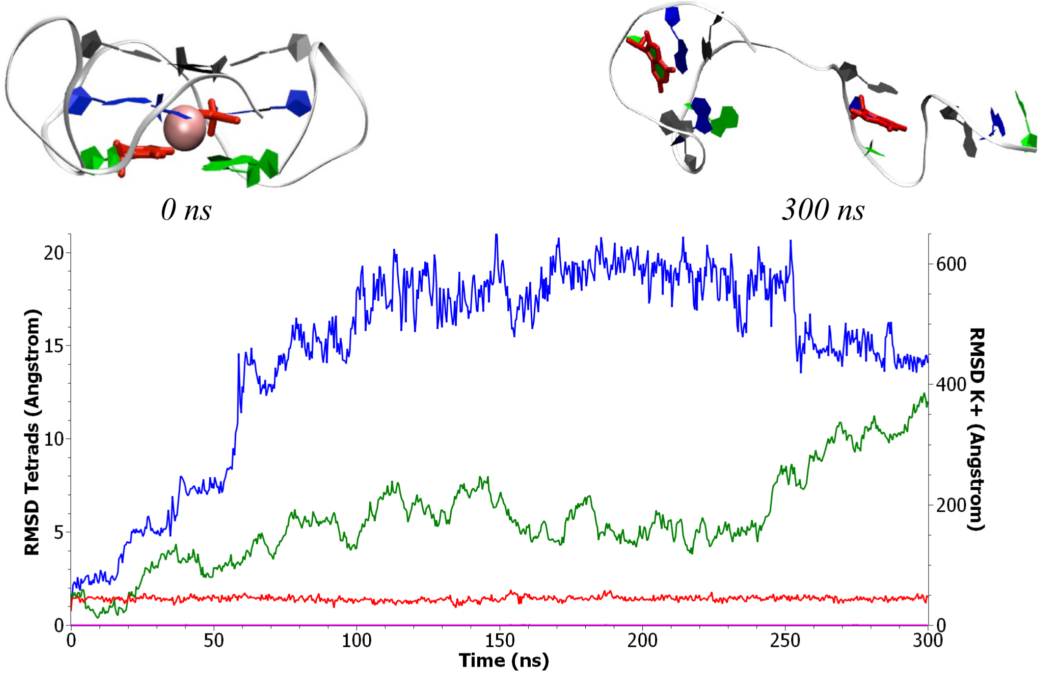


Run 1

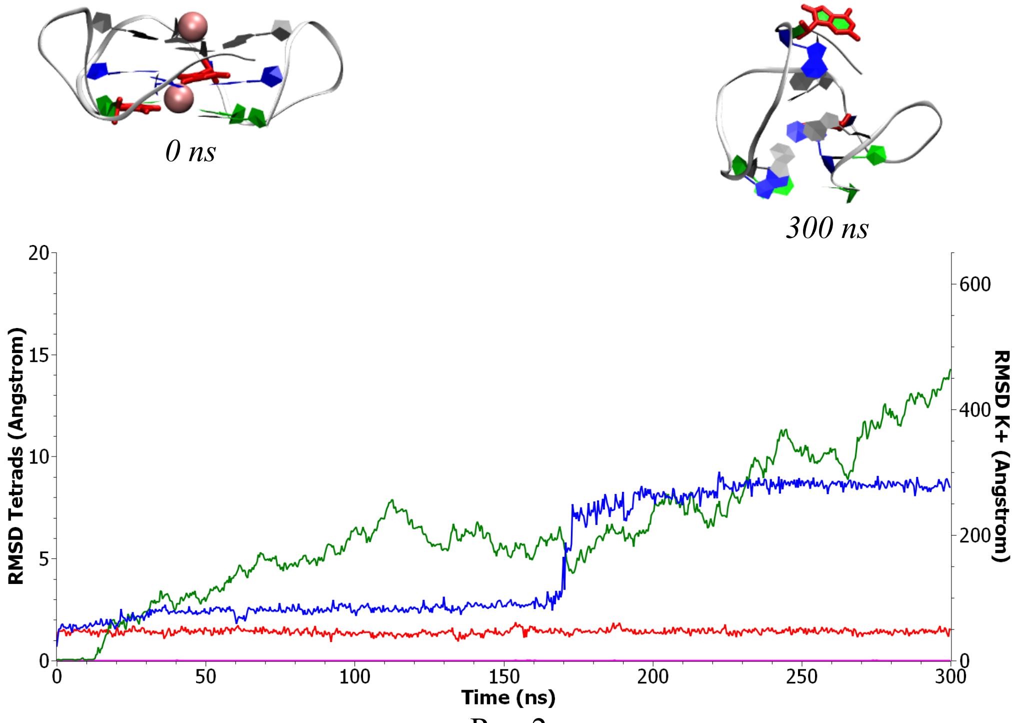


Run 2

Figure S16: Representative snapshots and time series of the RMSD for the two simulations involving parallel G4 presenting double lesions at position 4 and 15. *Red: RMSD of tetrads for the full undamaged structure. Blue: RMSD of tetrads for the full damaged structure. Pink: RMSD of $K^+$ for undamaged structure. Green: RMSD of $K^+$ for the damaged structure.*

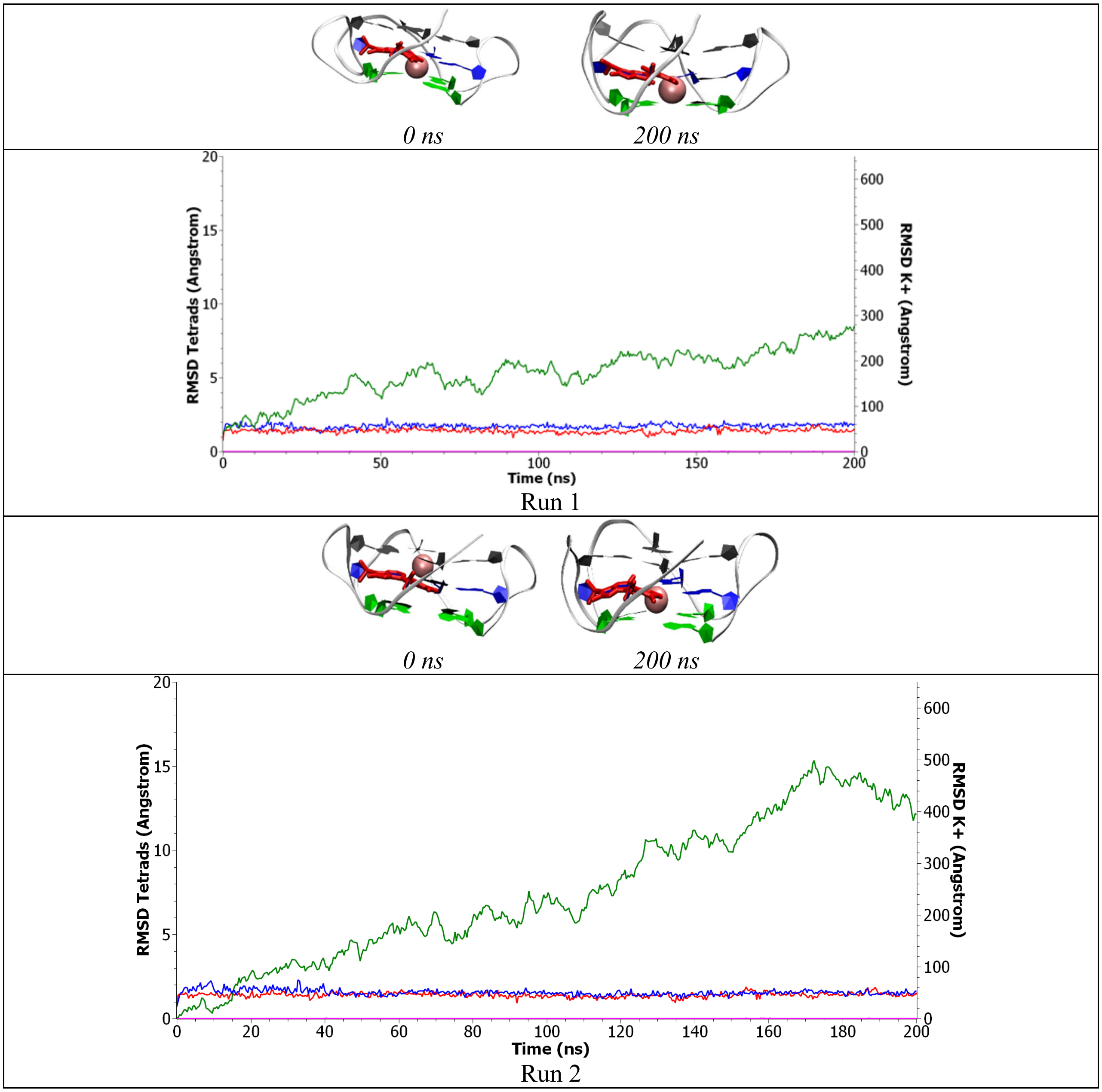


Figure S17: Representative snapshots and time series of the RMSD for the two simulations involving parallel G4 presenting double lesions at position 9 and 13. *Red: RMSD of tetrads for the full undamaged structure. Blue: RMSD of tetrads for the full damaged structure. Pink: RMSD of* $K^+$ *for undamaged structure. Green: RMSD of* $K^+$ *for the damaged structure.*

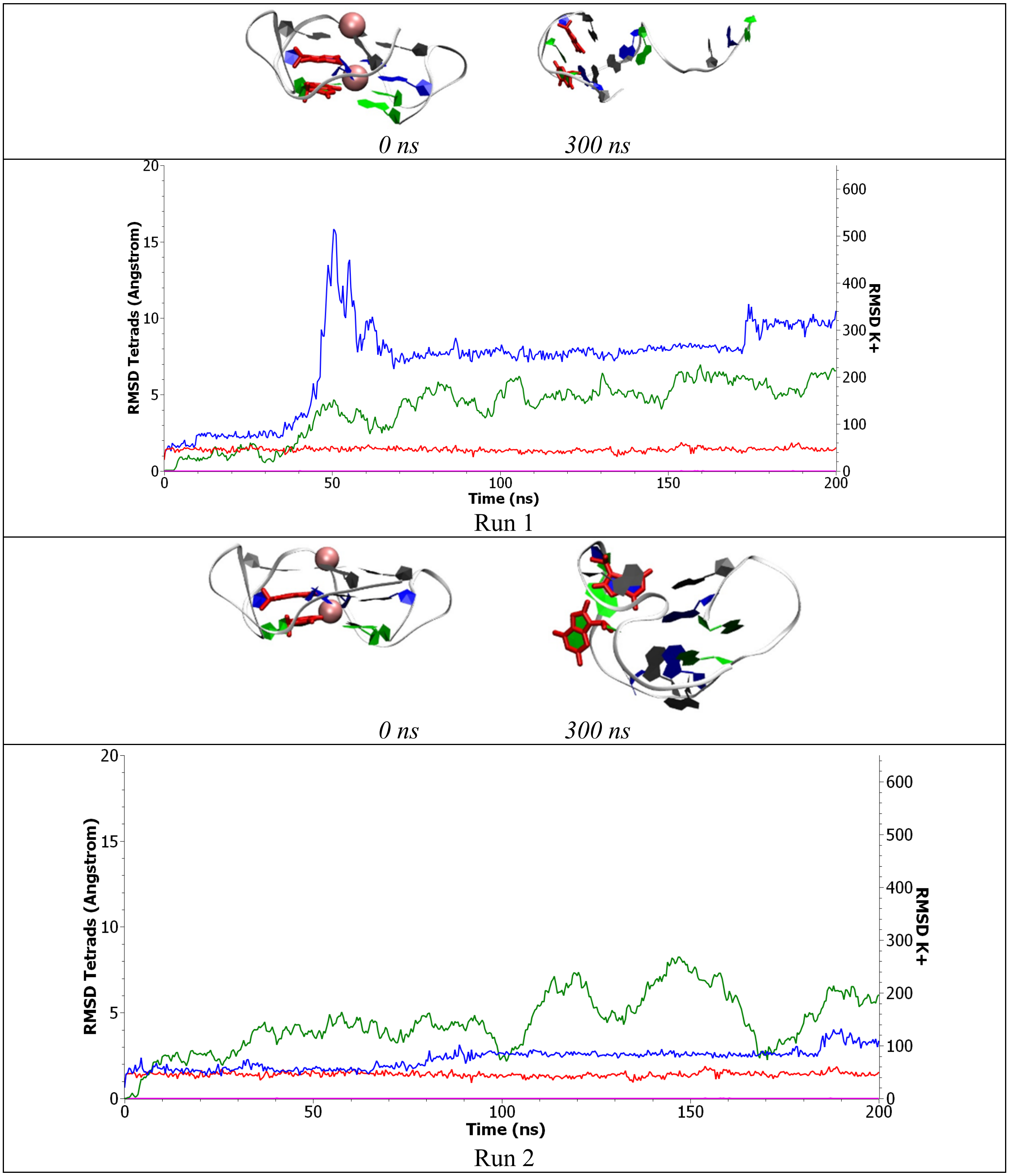


Figure S18: Representative snapshots and time series of the RMSD for the two simulations involving parallel G4 presenting double lesions at position 9 and 4. *Red: RMSD of tetrads for the full undamaged structure. Blue: RMSD of tetrads for the full damaged structure. Pink: RMSD of $K^+$ for undamaged structure. Green: RMSD of $K^+$ for the damaged structure.*

### Molecular dynamic simulations - Hybrid G4 - Single lesions

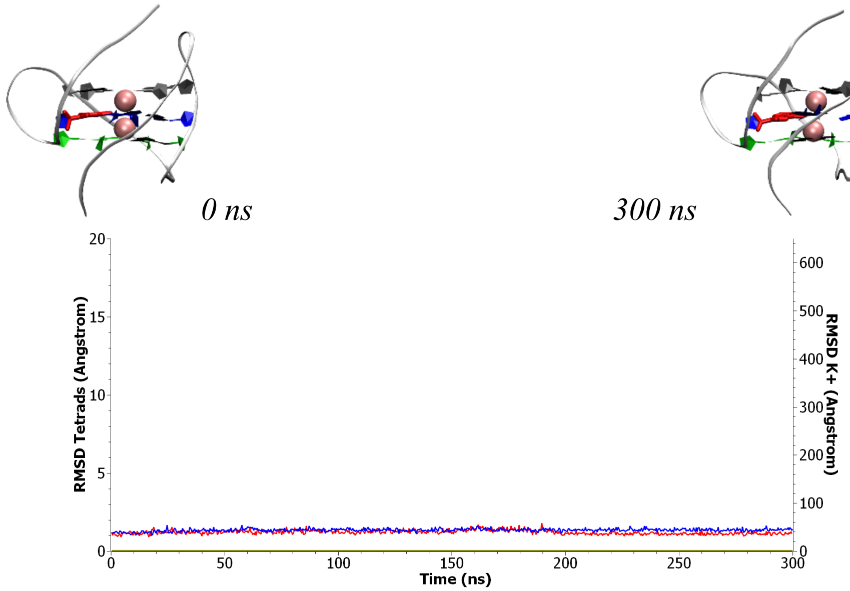


Run 1

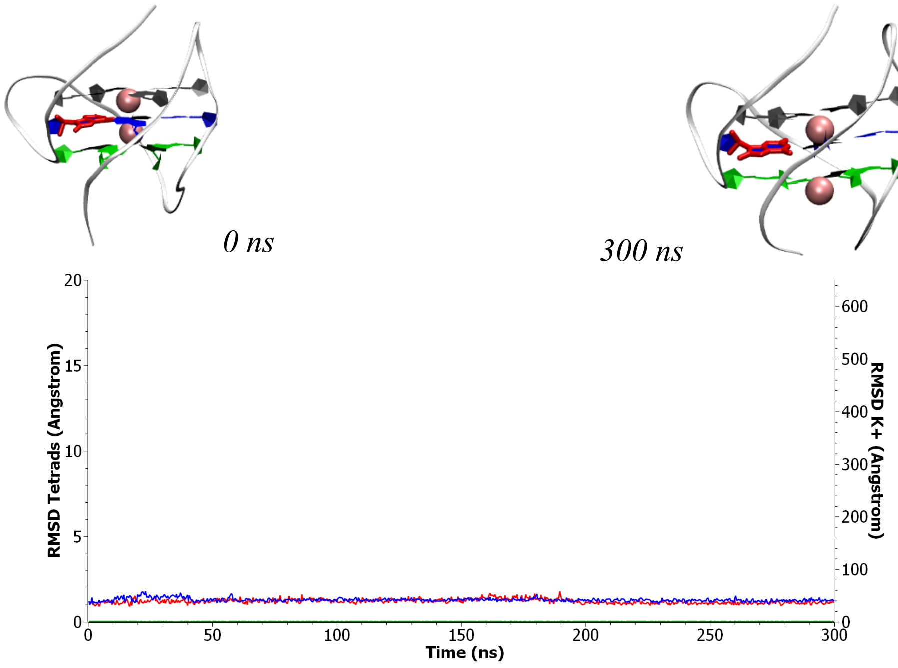


Run 2

Figure S19: Representative snapshots and time series of the RMSD for the two simulations involving hybrid G4 presenting a lesion at position 5. *Red: RMSD of tetrads for the full undamaged structure. Blue: RMSD of tetrads for the full damaged structure. Pink: RMSD of* $K^+$ *for undamaged structure. Green: RMSD of* $K^+$ *for the damaged structure.*

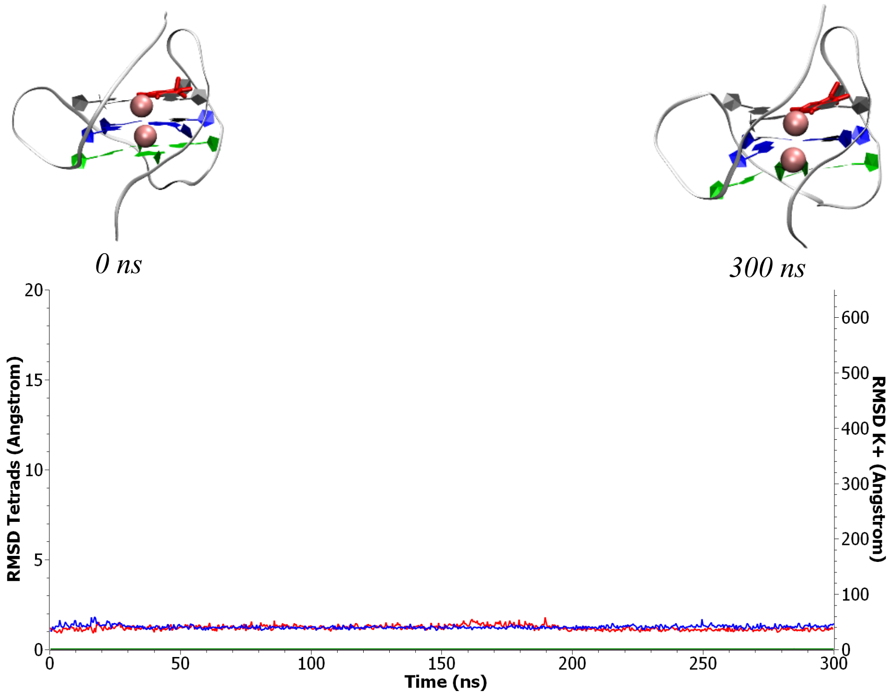


Run 1

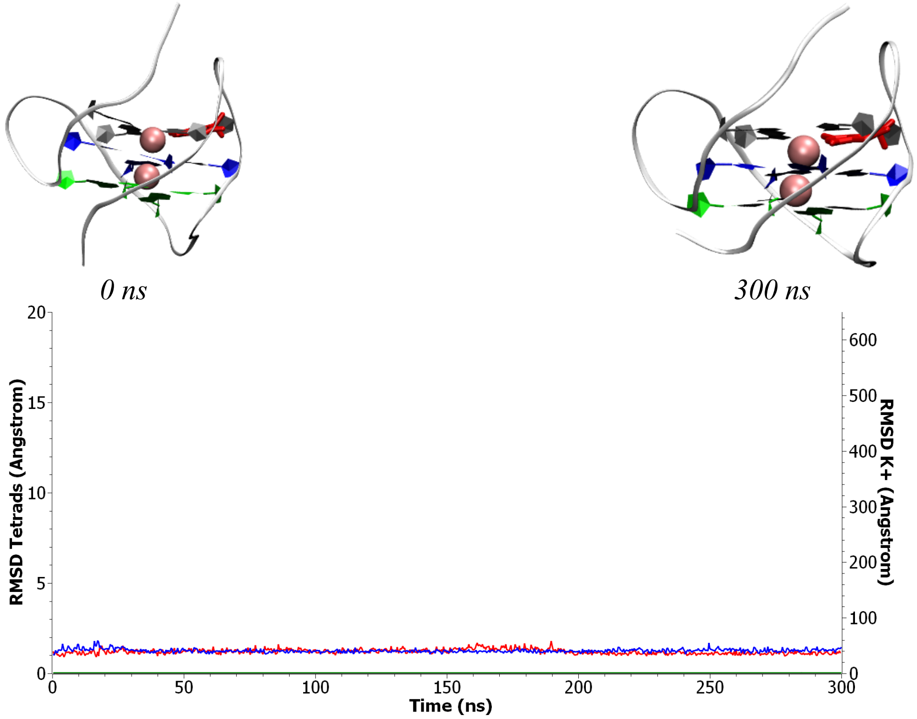


Run 2

Figure S20: Representative snapshots and time series of the RMSD for the two simulations involving hybrid G4 presenting a lesion at position 18. *Red: RMSD of tetrads for the full undamaged structure. Blue: RMSD of tetrads for the full damaged structure. Pink: RMSD of $K^+$ for undamaged structure. Green: RMSD of $K^+$ for the damaged structure.*

### Molecular dynamic simulations - Hybrid G4 - Double lesions

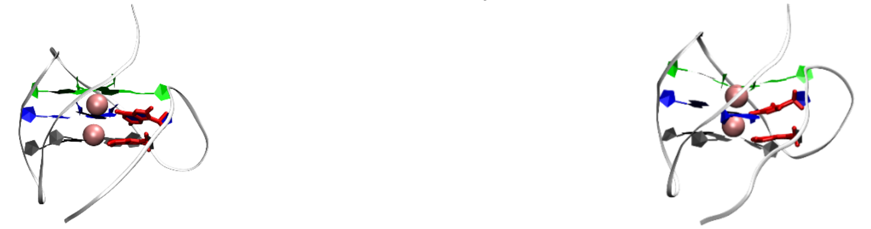

*0 ns* *300 ns*

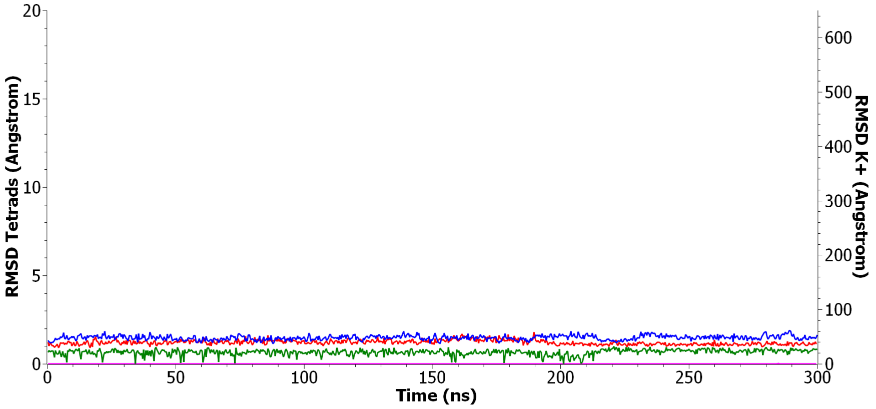


Run 1

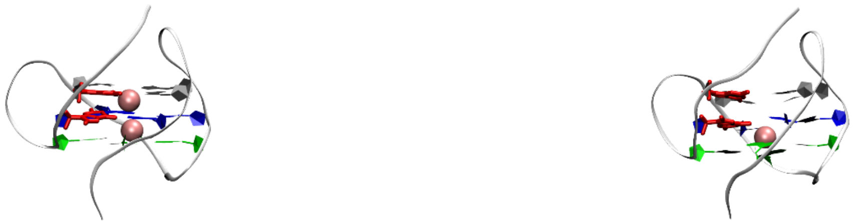

*0 ns* *300 ns*

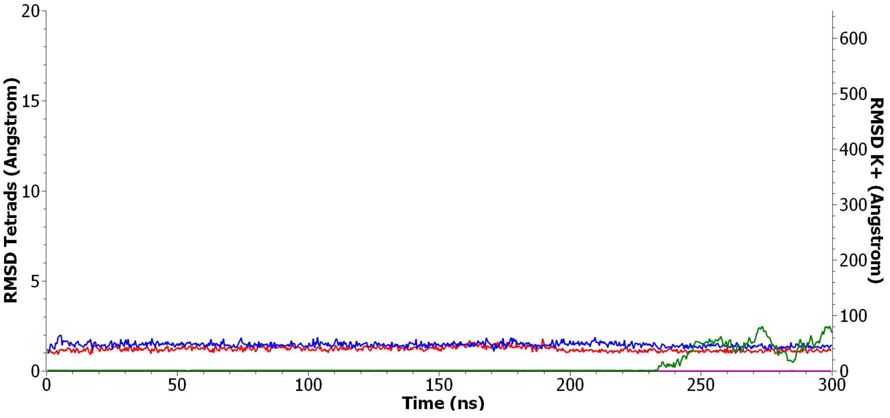


Run 2

Figure S21: Representative snapshots and time series of the RMSD for the two simulations involving hybrid G4 presenting double lesions at position 4 and 5. *Red: RMSD of tetrads for the full undamaged structure. Blue: RMSD of tetrads for the full damaged structure. Pink: RMSD of $K^+$ for undamaged structure. Green: RMSD of $K^+$ for the damaged structure.*

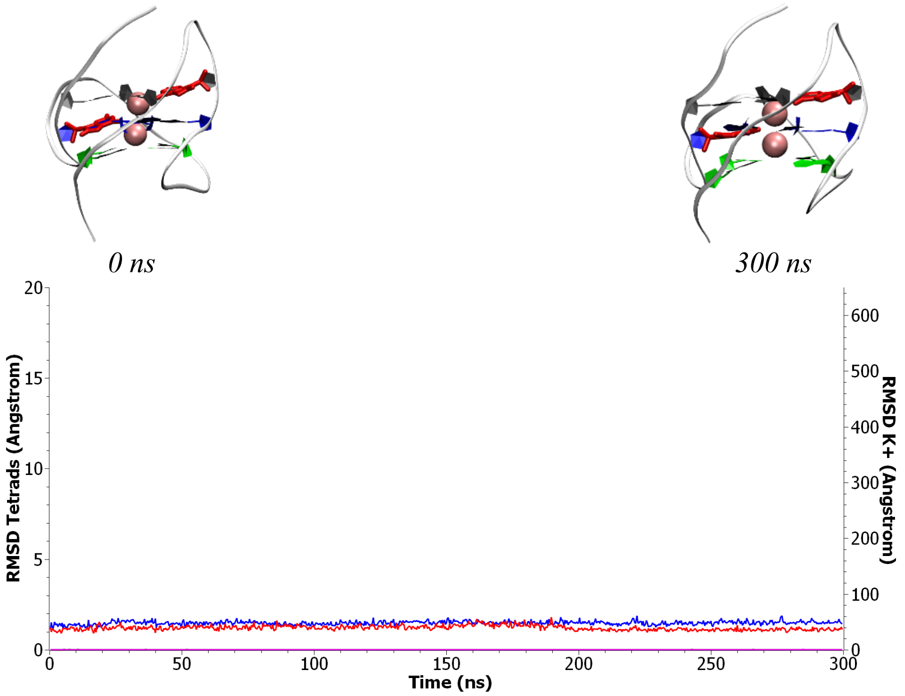


Run 1

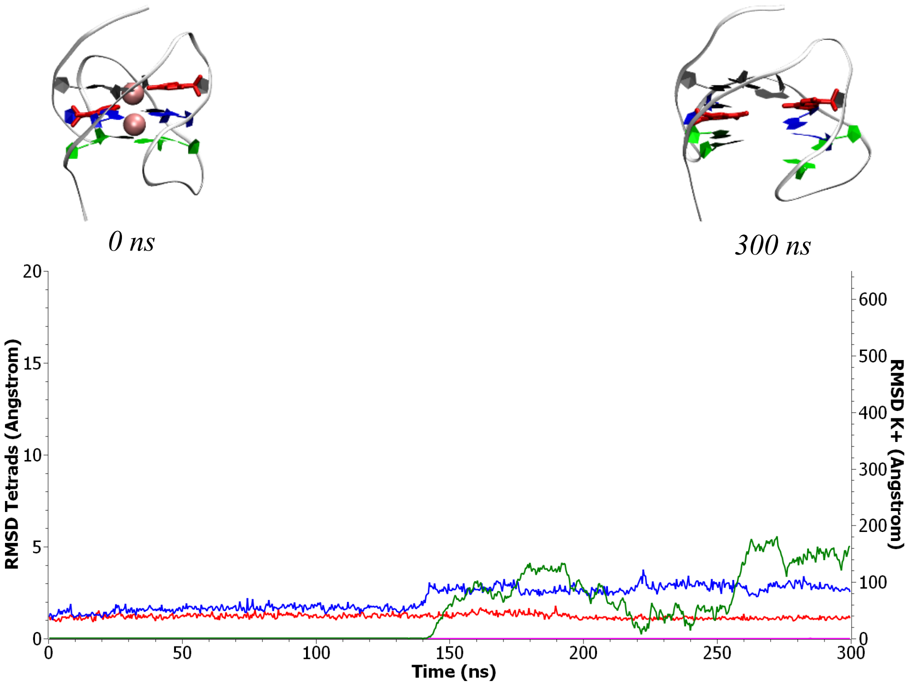


Run 2

Figure S22: Representative snapshots and time series of the RMSD for the two simulations involving hybrid G4 presenting double lesions at position 18 and 5. *Red: RMSD of tetrads for the full undamaged structure. Blue: RMSD of tetrads for the full damaged structure. Pink: RMSD of $K^+$ for undamaged structure. Green: RMSD of $K^+$ for the damaged structure.*

#### *Spectroscopic Titration*

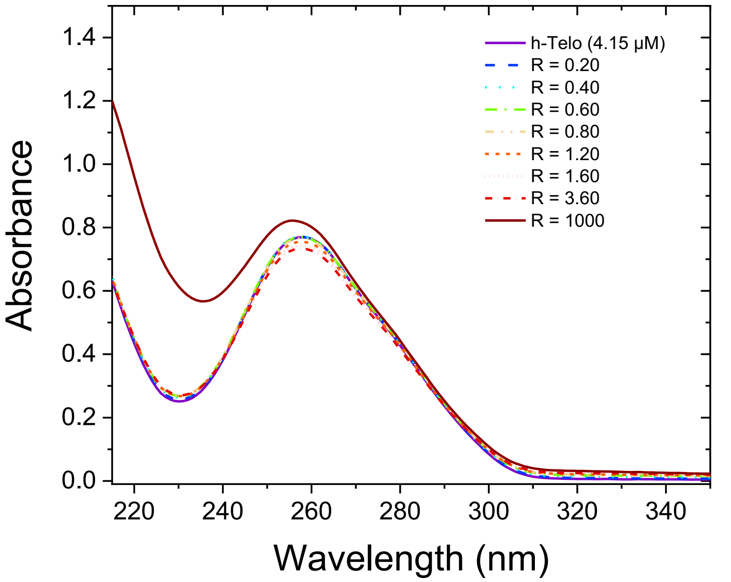


Figure S23: UV spectrum for hybrid G4 treated with different concentrations of $H_2O_2$

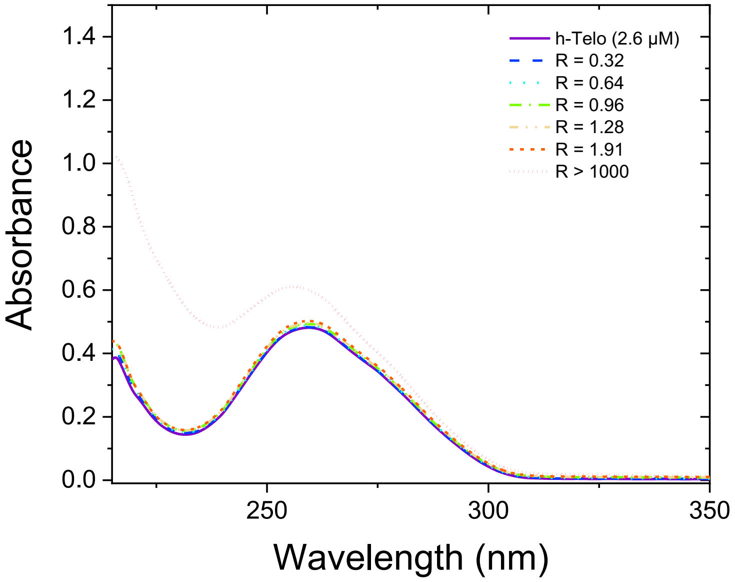


Figure S24: UV spectrum for hybrid G4 treated with different concentrations of $H_2O_2$

### *In cellulo* immunofluorescence - Microscopic photographs

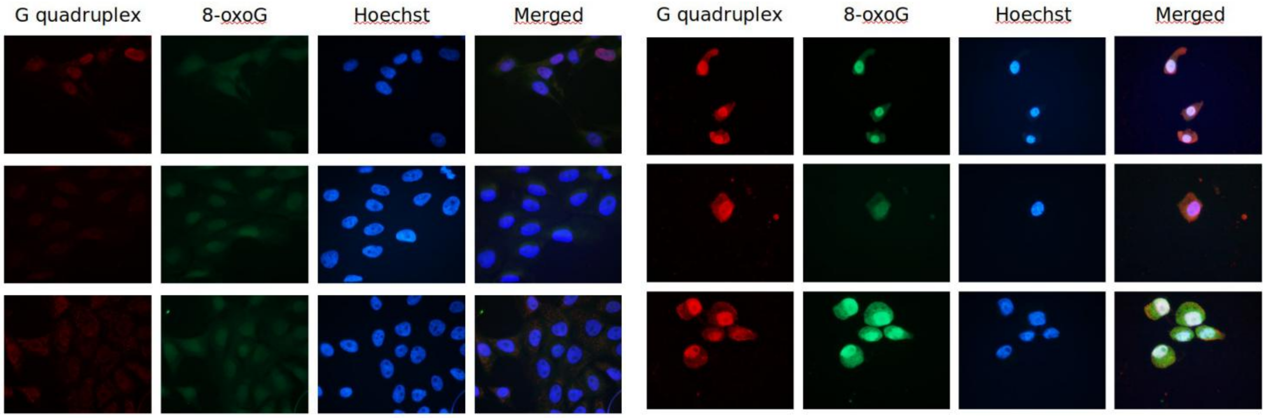


(a) Untreated cells. (b) Cells treated with 200 μM of $H_2O_2$ during 1h.

Figure S25: Immunofluorescence assay for MCF10a cells 200311

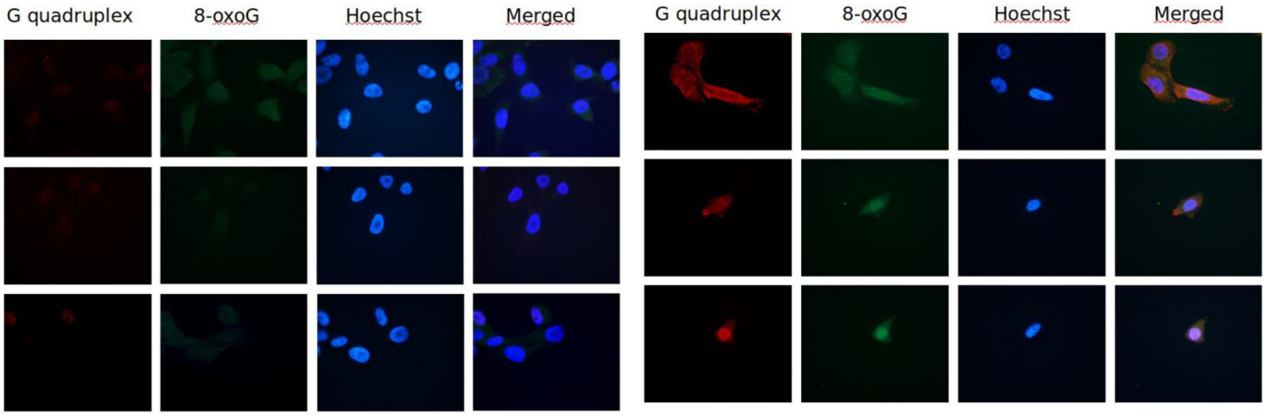


(a) Untreated cells. (b) Cells treated with 200 μM of $H_2O_2$, during 1 h.

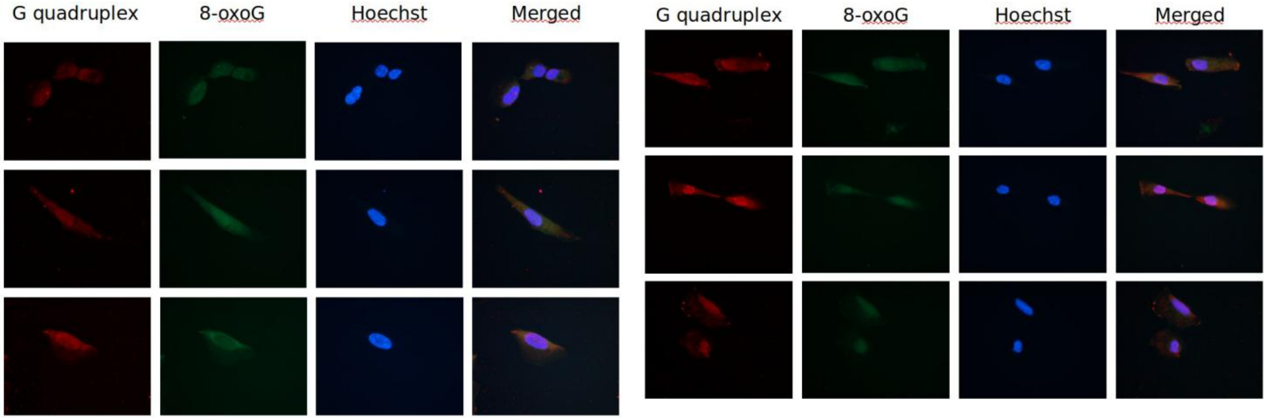


(c) Cells treated with 200 μM of $H_2O_2$ and 50 μM of ebselen, during 1 h. (d) Cells treated with 200 μM of $H_2O_2$ and 3 mM of tempol, during 1 h.

Figure S26: Immunofluorescence assay for MCF10a cells 200603 in presence of antioxidant

### *In cellulo* immunofluorescence - Quantification

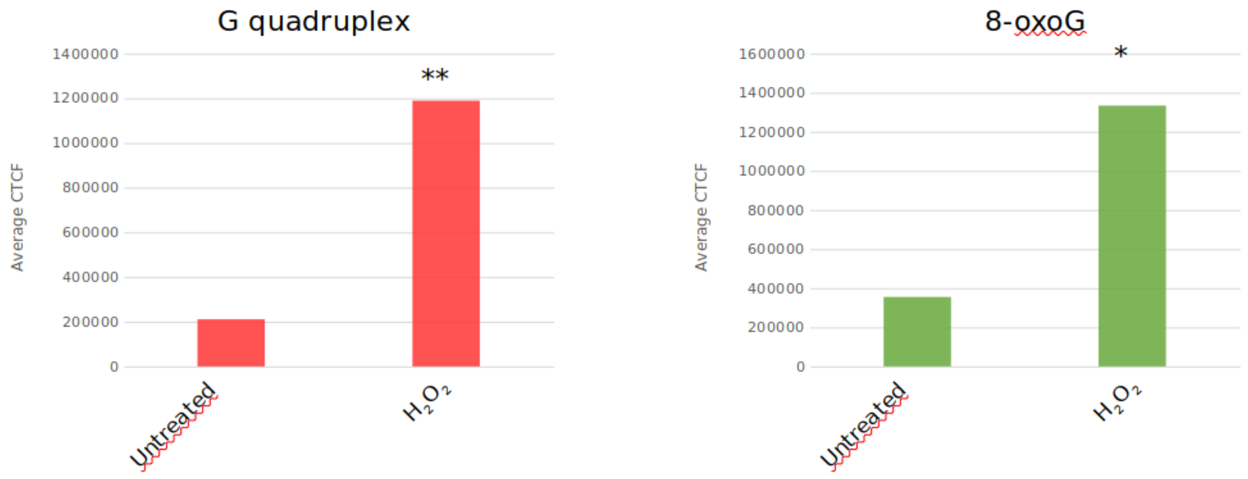


Figure S26: **MCF10a cells 200311.** *$H_2O_2$ : 200 µM for 1 h. T-test : * p<0,05, ** p<0,01, *** p<0,001.*

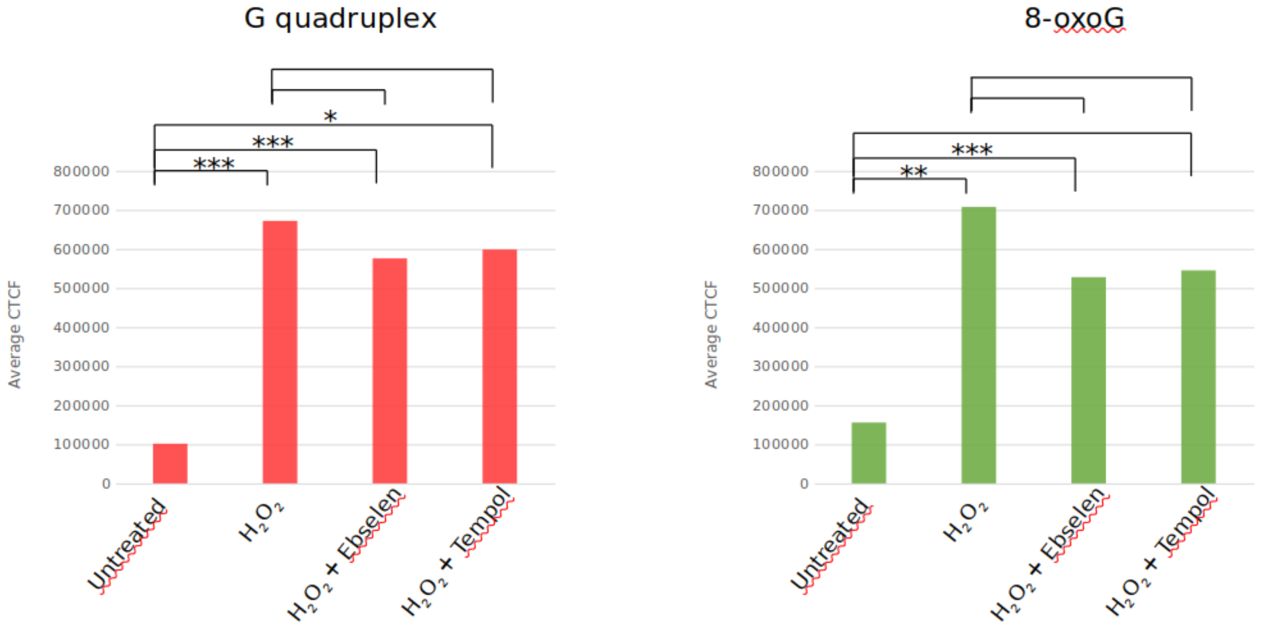


Figure S28: **MCF10a cells 200611.** *$H_2O_2$ : 200 µM $H_2O_2$ for 1 h. $H_2O_2$ + Ebselen : 200 µM $H_2O_2$ + 50 µM ebselen for 1 h. $H_2O_2$ + Tempol : 200 µM $H_2O_2$ + 3 mM tempol for 1 h. T-test : * p<0,05, ** p<0,01, *** p<0,001.*
